# Ca^2+^-activated CKL3 phosphorylates nucleoporin 58 to reprogram nuclear transport and execute effector-triggered immunity

**DOI:** 10.64898/2026.06.09.731143

**Authors:** Xing Zhang, Andres V. Reyes, Sargis Karapetyan, Yucong Xie, Yezi Xiang, Shou-Ling Xu, Xinnian Dong

**Affiliations:** Howard Hughes Medical Institute, Duke University, Durham, NC 27708, USA; Department of Biology, Duke University, Durham, NC 27708, USA; Department of Plant Biology and Carnegie Mass Spectrometry Facility, Carnegie Institution for Science, Stanford, CA 94305, USA

**Keywords:** effector-triggered immunity, Ca^2+^ influx, TurboID proximity labeling, nucleoporin 58, selective nuclear pore permeability, casein kinase 1-like 3

## Abstract

Recognition of pathogen effectors by nucleotide-binding leucine-rich repeat receptors (NLRs) activates effector-triggered immunity (ETI) in plants through resistosome formation, which generates a sustained cytosolic Ca^2+^ influx and ultimately leads to programmed cell death (PCD) at infection sites and resistance. However, the mechanisms linking Ca^2+^ influx to ETI execution remains a major knowledge gap. Using TurboID proximity labeling with the central transport channel nucleoporin 58 (Nup58) as a probe, we show that, instead of deregulation, ETI induction restricts general nuclear trafficking while selectively enhancing nuclear import of defense-related proteins. This switch is driven by elevated cytosolic Ca^2+^, which binds to the conserved Asp-149 in CASEIN KINASE 1-LIKE 3 (CKL3), promoting its interaction with and phosphorylation of Nup58 at Ser-149, thereby altering nuclear pore selectivity. This study identifies CKL3 and Nup58 as key regulators of ETI by establishing a mechanistic link from resistosome-mediated Ca^2+^ influx to nuclear transport reprogramming and immune execution.

## INTRODUCTION

Recent advances have elucidated key molecular mechanisms by which nucleotide-binding leucine-rich repeat receptors (NLRs), first identified over three decades ago,^1,2^ activate effector-triggered immunity (ETI) in plants. Upon recognition of pathogen effectors, both coiled-coil NLRs (CNLs) and Toll/interleukin-1 receptor NLRs (TNLs) oligomerize into resistosomes.^3–8^ CNL resistosomes function as Ca^2+^-permeable channels,^9^ whereas TNL resistosomes attain NADase activity, producing secondary messengers that activate helper NLRs which also form Ca^2+^-permeable channels.^10–14^ The resulting sustained Ca^2+^ influx is a central trigger of ETI signaling.^7–9,12^

Although ETI culminates in localized programmed cell death (PCD) and pathogen resistance, it is preceded by extensive transcriptomic and translatomic reprogramming,^15,16^ indicating that ETI is executed through a highly coordinated signaling cascade rather than a simple destructive process. Notably, both CNL- and helper NLR-resistosomes are presumed to localize predominantly to the plasma membrane,^9,12,17–19^ raising the fundamental questions: how does NLR-mediated Ca^2+^ influx at the cell periphery lead to transcriptomic changes in the nucleus and to what extent are these nuclear responses required for the execution of ETI?

The spatial separation between plasma membrane-localized immune signaling and nuclear transcriptional responses suggests a possible role of the nuclear pore complex (NPC) in regulating ETI. Indeed, the NPC has been implicated in plant immunity as mutants of the NPC components showed altered response to pathogens.^20–26^ Among them, CONSTITUTIVE EXPRESSION OF PATHOGENESIS-RELATED GENES 5 (CPR5) was found to be associated with nucleoporin 155 (Nup155) and its mutant exhibited autoimmune phenotype as well as spontaneous cell death, while CPR5 overexpression resulted in cytosolic retention of many nucleus-localized proteins, including ABSCISIC ACID INSENSITIVE 5 (ABI5) and NONEXPRESSOR OF PATHOGENESIS-RELATED GENES 1 (NPR1).^24^ However, these early reports did not address the questions whether and how NLR-triggered Ca^2+^ influx regulates the NPC permeability during ETI.

To identify the missing link between NLR-activated Ca^2+^ influx and downstream execution of ETI and to investigate a possible regulatory role that the NPC may play in the downstream signaling cascade, we employed a generic cargo protein to monitor nuclear transport activity during ETI, and applied TurboID-mediated proximity labeling using Nup58, a central channel nucleoporin known to regulate the selectivity of nucleocytoplasmic transport,^27,28^ to capture the ETI-regulated cargos. Surprisingly, instead of an overall deregulation of nuclear transport, we discovered that during ETI, the nuclear transport activity is enhanced selectively for certain defense-related proteins, while the general nuclear import activity is diminished. We further show that this change in the NPC’s selective permeability is independent of ETI-induced cargo gene transcription and translation. Instead, it is triggered upon phosphorylation of Nup58 at Ser-149 carried out by CASEIN KINASE 1-LIKE 3 (CKL3) whose interaction with Nup58 is enhanced through association with Ca^2+^ during ETI. Consequently, the change in NPC selectivity leads to transcriptional reprogramming required for ultimate PCD and resistance. Therefore, this study establishes a mechanistic link between NLR-mediated Ca^2+^ influx and PCD execution by positioning CKL3 and Nup58 as key intermediate regulators in the ETI signaling pathway.

## RESULTS

### NLR-mediated Ca^2+^ influx affects nuclear import activity of the NPC

To fill the knowledge gap in the signaling cascade from resistosome-mediated Ca^2+^ influx in the cytosol to transcriptomic reprogramming in the nucleus, we examined changes in the nuclear transport activity during ETI, using YFP and luciferase (LUC) as inert cargo reporters without disturbing the endogenous biological functions, instead of transcriptional regulators used in previous studies.^24^ As expected, a single YFP (∼ 27 kDa) is freely diffusible through the nuclear pore, exhibiting both nuclear and cytosolic localization when transiently expressed in *Nicotiana benthamiana* leaves, while the 2×YFP-LUC fusion protein (∼ 116 kDa) is too large to be passively diffused through the nuclear pore and enter the nucleus (Figures S1A and S1B). However, addition of a nuclear localization sequence (NLS) to 2×YFP-LUC resulted in its facilitated import into the nucleus (Figure S1C), thereby making 2×YFP^NLS^-LUC a generic, inert cargo reporter to monitor nuclear transport activity during ETI.

To establish a transient ETI-induction assay in *N. benthamiana*, we leveraged the knowledge that the bacterial effectors AvrRpt2 and AvrRpm1 activate ETI in *Arabidopsis* by cleaving or ADP-ribosylating the key immune repressor protein RPM1-INTERACTING PROTEIN 4 (RIN4), whose integrity is normally under the surveillance of the CNL receptor RESISTANCE TO PSEUDOMONAS SYRINGAE 2 (RPS2) and RESISTANCE TO PSEUDOMONAS SYRINGAE PV. MACULICOLA 1 (RPM1), respectively.^17,29–33^ Because of the polymorphisms in NbRIN4, expressing AtRPS2 or AtRPM1 in *N. benthamiana* alone is sufficient to trigger ETI-associated PCD.^34,35^ We first tagged CFP to the C-terminus of RPS2 and put its transcription under the control of dexamethasone (Dex). After expressing this fusion construct in *N. benthamiana* leaves, we detected Dex-induced PCD in a dose-dependent manner, using both cell death-accompanied leakage of ions quantified through conductivity, and cell death-associated autofluorescence imaged with Infrared Fluorescence (IR800) (Figures 1A and 1B). In contrast, when CFP was fused to the N-terminus of RPS2, cell death did not occur (Figures 1A and 1B), similar to the failure of N-terminal tagged ZAR1 in inducing cell death.^9^

**Figure 1.**
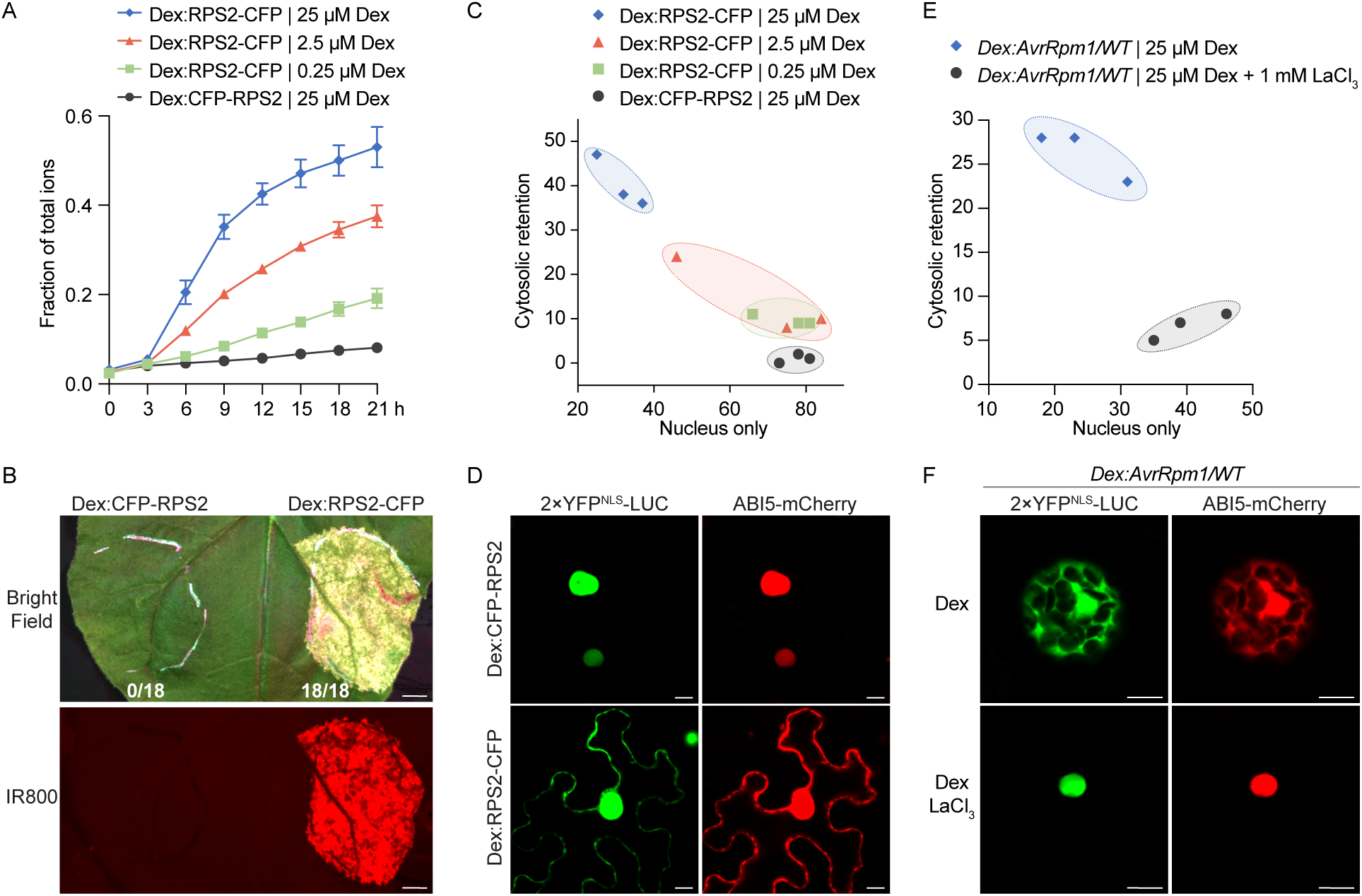
NLR-mediated Ca^2+^ influx affects nuclear import activity of the NPC. (A and B) RPS2-induced cell death in *Nicotiana benthamiana*. Leaves were inoculated with *Agrobacterium* carrying the dexamethasone-inducible *RPS2-CFP* (*Dex:RPS2-CFP*) or the inactive *Dex:CFP-RPS2* and incubated for 36 h. RPS2-induced plant cell death was assessed following Dex induction (0.25 μM, 2.5 μM, and 25 μM) in a time-course conductivity assay, measuring ion leakage normalized to the total ion content (A), and by counting dead versus total inoculated leaves as well as by using Infrared Fluorescence imaging (IR800) to visualize cell death-associated autofluorescence with images captured 1 day after Dex treatment (25 μM) (B). Data are presented as means ± SEM (n = 3) per timepoint for (A) and scale bar = 0.5 cm for (B). (C and D) RPS2-triggered cytosolic retention of 2×YFP^NLS^-LUC and ABI5-mCherry in *N. benthamiana*. Leaves were co-inoculated with *Agrobacteria* carrying *Dex:RPS2-CFP* or the inactive *Dex:CFP-RPS2* together with *35S:2*×*YFP^NLS^-LUC* and *35S:ABI5-mCherry* as in (A). Confocal images were taken 3 h after Dex treatment (0.25 μM, 2.5 μM, and 25 μM) and cytosolic retention of 2×YFP^NLS^-LUC was quantified from three independent repeats with ∼100 cells counted per repeat (C). Representative cell images with cytosolic retention of 2×YFP^NLS^-LUC and ABI5-mCherry upon Dex treatment (25 μM) are shown in (D). Scale bar = 10 μm. (E and F) AvrRpm1-induced, Ca^2+^-dependent cytosolic retention of 2×YFP^NLS^-LUC and ABI5-mCherry in *Arabidopsis* protoplasts. Protoplasts derived from *Dex:AvrRpm1/WT* transgenic plants were transfected with *35S:2*×*YFP^NLS^-LUC* and *35S:ABI5-mCherry*. Images were taken 2 h after treatment with 25 μM Dex or Dex plus 1 mM LaCl_3_ with the cytosolic retention of 2×YFP^NLS^-LUC quantified from three independent repeats (∼70 protoplasts per repeat) (E). Representative cell images with cytosolic retention of 2×YFP^NLS^-LUC and ABI5-mCherry upon Dex treatment (25 μM) are shown in (F). Scale bar = 10 μm.

During this RPS2-induced cell death in *N. benthamiana*, we observed a Dex dose-dependent cytosolic retention of 2×YFP^NLS^-LUC (Figure 1C). A similar phenomenon was observed for the ABI5-mCherry protein, which also contains an NLS and normally undergoes facilitated nuclear import (Figures 1C and 1D). In addition to RPS2, we also performed Dex-induced expression of CNL RPM1 and the autoactive variant (D485V) of the helper NLR N REQUIREMENT GENE 1 (NRG1.DV)^12^ and observed similar cytosolic retention of 2×YFP^NLS^-LUC as well as cell death (Figures S1D and S1E). These findings indicate that a constraint on nuclear localization is a shared downstream response upon NLR activation.

Finally, to determine whether these ETI-induced changes in the NPC permeability are Ca^2+^-dependent, we used the *Arabidopsis* protoplasts generated from the transgenic plants carrying the Dex-inducible bacterial effector AvrRpm1 (*Dex:AvrRpm1/WT*). We found that when the Ca^2+^ influx was blocked by lanthanum chloride (LaCl_3_) treatment, cytosolic retention of 2×YFP^NLS^-LUC and ABI5-mCherry induced by AvrRpm1 was inhibited (Figures 1E and 1F), indicating that the NLR-mediated change in NPC permeability requires Ca^2+^ influx.

### Tb-Nup58 captures cargos passing through the nuclear pore during ETI

The observed constraint on nuclear localization during ETI appears to conflict with the transcriptional reprogramming which requires transcription factors (TFs) to be in the nucleus. To capture the cargos transported through the nuclear pore during ETI *in planta*, we performed a TurboID proximity labeling experiment focusing on Nup58, which is a phenylalanine-glycine (FG) repeat-containing channel Nup in the center of the NPC (Figure 2A). We generated a transgenic line expressing both TurboID-3×HA-Nup58 (Tb-Nup58) and Dex-inducible AvrRpt2 (*Dex:AvrRpt2*) for a time-course induction of ETI. We chose the time points prior to the onset of PCD, based on the ion leakage measurement, to capture the possible early ETI/PCD regulators (Figures S2A and S2B). For calibration, we used the transgenic line expressing 2×YFP-TurboID (YFP), which has been shown in multiple previous TurboID experiments to be ubiquitously distributed in *Arabidopsis* cells.^36^ Principal Component Analysis (PCA) showed a clear separation of the YFP control from both H_2_O- and Dex-treated Nup58 samples (Figure S2C), indicating the specificity of the targets captured by the two different baits. Based on the log_2_fold change (log_2_FC) of label free quantification (LFQ) intensity of all the Nup58 samples compared to the YFP control (log_2_FC > 1 and -log_10_ (p-value) > 1), we designated the 518 Nup58-captured proteins as the Nup58-proxiome (Figure S2D and Table S1).

**Figure 2.**
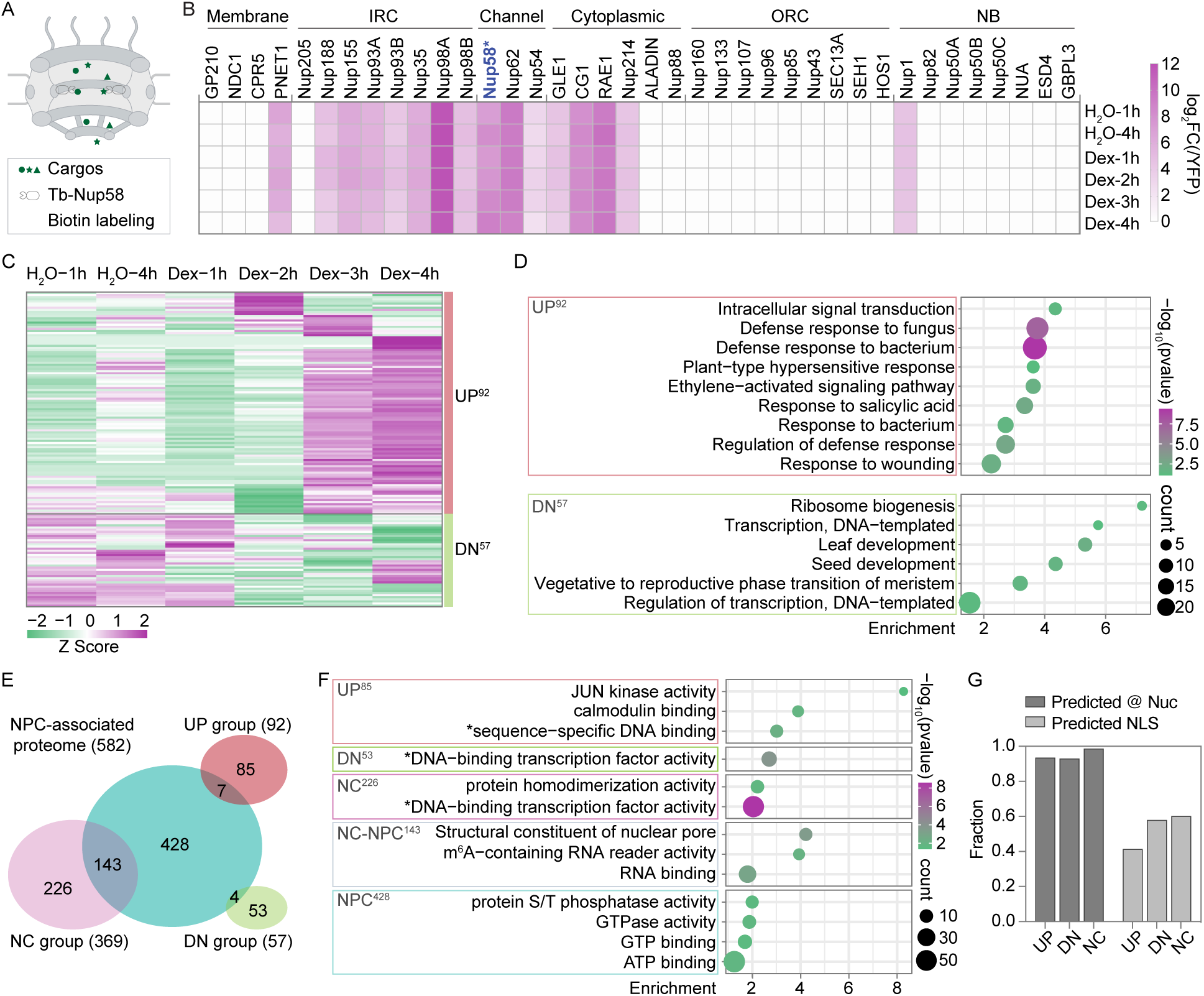
Tb-Nup58 captures cargos passing through the nuclear pore during ETI. (A) Schematic illustration of using TurboID-3×HA-Nup58 (Tb-Nup58) as a bait to capture the Nup58-proxiome. The *35S:Tb-Nup58* transgenic *Arabidopsis* line used in the experiment also carries *Dex:AvrRpt2* for ETI induction. (B) Heatmap depicting changes in nucleoporin levels captured by Tb-Nup58 with and without ETI induction triggered by Dex-mediated expression of *AvrRpt2*. Enrichment was quantified as log_2_fold change (log_2_FC) for each Tb-Nup58 sample relative to the corresponding 2×YFP-TurboID (YFP) control. IRC, inner ring complex; ORC, outer ring complex; NB, nuclear basket. (C) Clustered heatmap showing proteins with increased (UP) or decreased (DN) capture by Tb-Nup58 during ETI within the Nup58-proxiome. Superscript numbers indicate the number of proteins in each group. (D) Geno Ontology (GO) analysis of enriched biological processes in the UP- and the DN-cargo groups, using the Nup58-proxiome as the background. (E) Venn diagram of overlapping proteins between Nup58-proxiome groups (UP, DN, and NC) and the NPC-associated proteome reported by Tang et al.^37^ (F) GO term analysis of enriched molecular functions performed for cargo groups identified only in the Nup58-proxiome following ETI induction (UP^85^, DN^53^, and NC^2^^26^), as well as cargo groups shared with the NPC-associated proteome reported by Tang et al. (NC-NPC^1^^43^) and those specific to the Tang et al (NPC^4^^28^), using the combined protein pool as the background. *, Transcription-related GO terms. (G) Fractions of the Nup58-proxiome groups predicted to be localized in the nucleus (@ Nuc, SUBA5^43^) or to carry an NLS (LOCALIZER 1.0.4^44^).

We found enrichment of Nups that are known to be in close proximity to Nup58, including all channel Nups, the majority of Inner Ring Complex (IRC) Nups, and several cytoplasmic Nups (Figure 2B), indicating the success of the experiment. Interestingly, we did not observe any significant changes in the association between Nup58 and other Nups in response to Dex-induced ETI (Figure 2B), suggesting that the NPC remained as an intact unit during the detected time period of ETI response. We then performed Gene Ontology (GO) enrichment analysis for the Nup58-proxiome. We found the “structural constituent of the nuclear pore” as the most enriched Molecular Function (MF), together with several nucleus-related processes, including transcription, RNA processing, protein and RNA transporting within the enriched terms in MF and Biological Processes (BP) (Figure S2E), further validating the robustness of our TurboID experiment. Notably, while the enrichment of proteins from most BP terms was unaffected during ETI based on their LFQ intensities, the proteins involved in “cellular response to hypoxia” and “regulation of defense response” were preferentially enriched upon Dex treatment (Figures S2E and S2F).

To classify the targets based on their enrichment patterns upon ETI induction, we generated a clustered heatmap of the Nup58-proxiome and designated the 92 proteins with ETI-enhanced capture as the UP^92^ group, while the 57 proteins with reduced capture as the down (DN^57^) group (Figure 2C). Consistent with the results in Figure S2F, defense responses were highly enriched BPs in the UP group, even with the Nup58-proxiome as the background for the analysis, whereas development-related terms were enriched in the DN group (Figure 2D), suggesting that, instead of deregulation, a preferential cargo transport is triggered during ETI.

Recently, an NPC-associated proteome was reported, showing enrichment of chromatin remodelers, transcriptional regulators, and mRNA processing machinery on the nucleoplasmic side of the NPC, and enrichment of translation regulatory machinery on the cytoplasmic side of the NPC.^37^ Surprisingly, comparing our Nup58-proxiome to the abovementioned NPC-associated proteome, we found that the majority of the ETI-induced UP and DN targets were specific to our Nup58-proxiome, whereas the shared targets predominantly fell into the non-changed (NC) category (Figure 2E, Table S2). We then performed GO term enrichment in MFs for each new category of Nup58-specific targets (UP^85^, DN^53^ and NC^226^), using the combined NPC-associated proteome and Nup58-proxiome as the background, and found that the enriched MFs across all three categories were predominantly associated with transcription. Moreover, the ETI-induced UP^85^ that is unique to the Nup58-proxiome was further enriched for proteins involved in calmodulin binding, such as TFs CBP60g and SARD1 involved in the biosynthesis of the immune signal salicylic acid (SA),^38–40^ as well as proteins exhibiting JUN kinase activity, including the known central immune signaling component MPK3^41,42^ (Figure 2F). These enrichments further demonstrate that our Nup58-proxiome effectively captured the dynamic changes in the transport activity of the NPC during ETI. Curiously, although most proteins in all three Nup58-proxiome groups are predicted to localize to the nucleus,^43,44^ roughly half carry a known NLS (Figure 2G).

### ETI enhances nuclear import of defense-related proteins while constraining general cargo transport

The enhanced capture of cargos in the UP group by Tb-Nup58 during ETI suggests a potential reprogramming, instead of deregulation, of nuclear transport activity. To identify the factors underlying this cargo preference during ETI, we first examined proteins’ molecular weight (MW) and isoelectric point as potential determinants. However, no significant differences were observed (Figures S3A and S3B). Given the highly correlated transcriptomic and translatomic changes during ETI,^15^ we then assessed the ETI-induced changes in transcript levels (RNA-seq, RS) and translational activity (ribosome footprinting, RF) for all genes encoding the Nup58-proxiome to examine a possible correlation between expression levels and cargo capture patterns. Consistently, highly correlated transcriptional (log_2_RSfc) and translational (log_2_RFfc) changes were observed for the overall Nup58-proxiome genes during ETI (Figures S3C-S3E). However, 47 of the 92 UP-group proteins exhibited no increase in transcriptional or translational regulation (Figure S3C), and 52 of the 57 DN-group proteins showed no corresponding downregulation (Figure S3D). These findings suggest that the transcriptional and/or translational changes are unlikely to be the primary drivers for the observed switch in the nuclear transport selectivity during ETI.

To test whether the TurboID-captured cargo transport changes reflect an alteration in the nuclear import activity, we examined the distribution of constitutively expressed, fluorescent-tagged cargos in the nucleoplasm (Nuc) versus in the cytoplasm (Cyt) with and without ETI induction (Figure 3A). We observed a decreased Nuc/Cyt ratio for the generic 2×YFP^NLS^-LUC reporter upon ETI triggered by Dex-induced RPS2 expression in *N. benthamiana* (Figure 3B), indicating a more constrained general nuclear import activity. Similar reductions in Nuc/Cyt ratios were observed for cargos from the DN group (Figures 3C, S3D and S3F). In contrast, we observed increased Nuc/Cyt ratios for the UP-group cargos regardless of whether their endogenous genes were induced (e.g., CBP60g, SARD1, and MPK3) or not (e.g., WRKY11, EIN3, and CAMBP25) during ETI (Figures 3D, 3E, S3C, S3G and S3H). These findings support our hypothesis that there is a distinct mechanism that modulates NPC permeability to preferentially import certain defense-related cargos into the nucleus, while simultaneously constraining the general nuclear import activity upon ETI induction.

**Figure 3.**
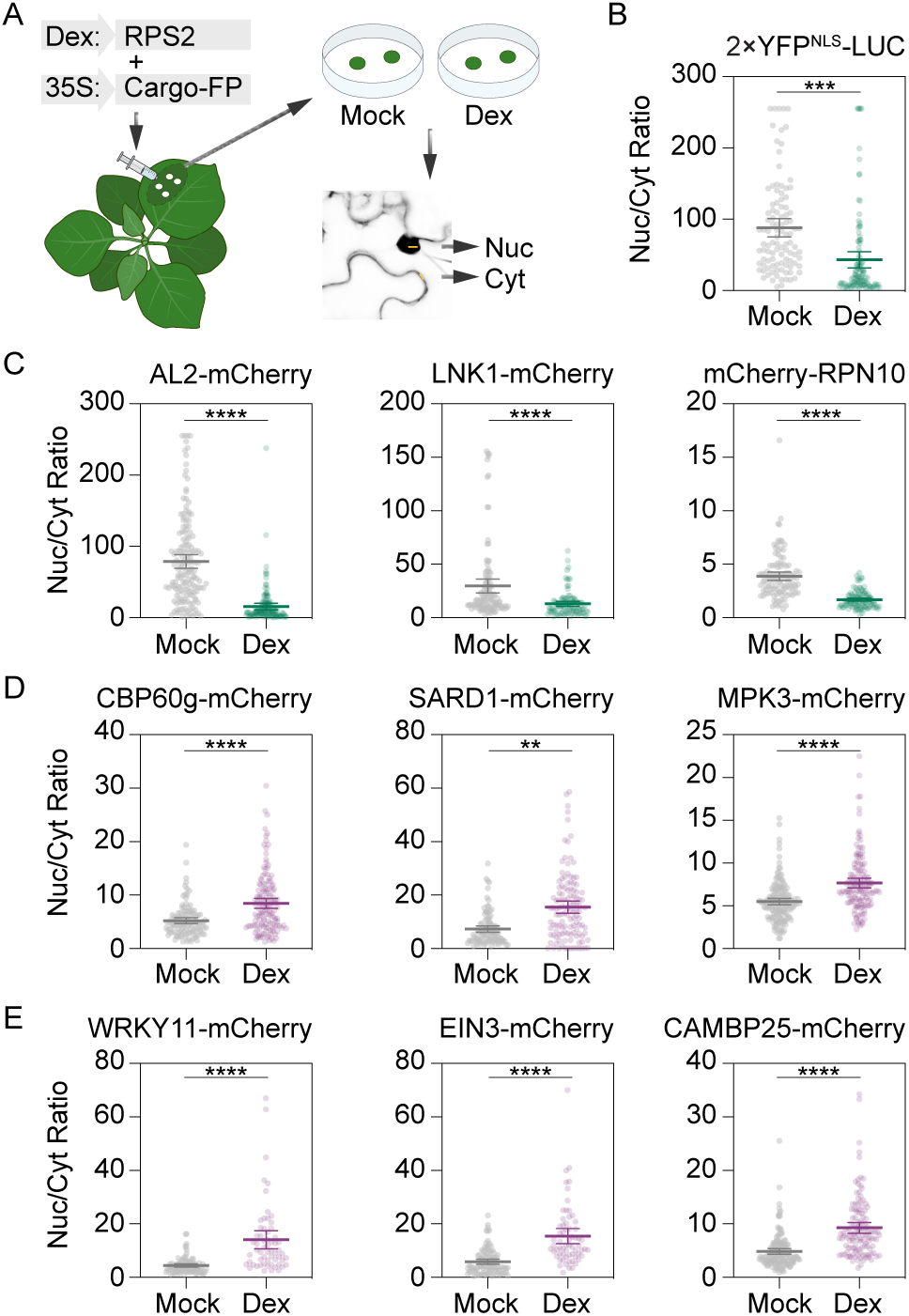
ETI enhances nuclear import of defense-related proteins while constraining general cargo transport. (A) Schematic illustration of monitoring nuclear transport of cargo proteins during ETI. *N. benthamiana* leaves were co-inoculated with *Agrobacteria* carrying *Dex-inducible RPS2* and *35S:Cargo-fluorescent protein* (*FP*) for 36 h. Confocal images were taken 3 h after water (Mock) or 25 μM Dex treatment. The fluorescence intensities in both the nucleus (Nuc) and the cytosol (Cyt) of the same cell were measured simultaneously, and the Nuc/Cyt ratio was then quantified. (B and C) Changes in nuclear import of the generic reporter 2×YFP^NLS^-LUC (B) and DN-group cargo-FP from the Nup58-proxiome (C) in response to ETI induction, measured as described in (A). (D and E) Changes in nuclear import of UP-group cargo-FP from the Nup58-proxiome in response to ETI induction, measured as described in (A). Representative cargos with and without transcriptional and translational induction of their endogenous genes (Figure S3C) are shown in (D) and (E), respectively. Each dot in the scatter plots represents a single cell, and the line represents mean ± 95% CI. Individual columns were compared using Student’s *t*-test. **p < 0.01, ***p<0.001, ****p<0.0001.

### Phosphorylation of Nup58 at Ser-149 is required for the selective nuclear import of defense proteins and ETI

To reveal the mechanism of the ETI-induced switch in selective nuclear import, we re-examined our TurboID LC-MS/MS data to search for phosphorylation in Nups, since this modification and/or protein stability have been reported to regulate Nups during the nuclear envelop breakdown associated with cell division or permeability change upon stress in animals.^45–52^ Notably, we detected two ETI-specific phospho-peptides from Nup58 with five candidate serine residues (Figure S4A, Table S3). When expressed in *N. benthamiana*, we observed an upshifted HA-Nup58 band on a Phos-tag gel upon Dex-induced RPS2 expression (Figure 4A), supporting the phosphorylation of the protein detected in LC-MS/MS. Among the ETI-specific phospho-peptides, only phosphorylation at Ser-149 showed clear and reliable MS2 spectra (Figures S4B-S4D). Consistently, when we generated the phospho-mimic variant for each candidate serine residue in Nup58, we found that only the variant HA-Nup58^S149D^ led to cytosolic retention of the 2×YFP^NLS^-LUC reporter and PCD when transiently expressed in *N. benthamiana* (Figures 4B, S4E and S4F), suggesting that Ser-149 is the ETI-induced phosphorylation site in Nup58. Further supporting this conclusion, the phospho-dead HA-Nup58^S149A^ variant diminished the RPS2-induced band shift on the Phos-tag gel (Figure 4C).

**Figure 4.**
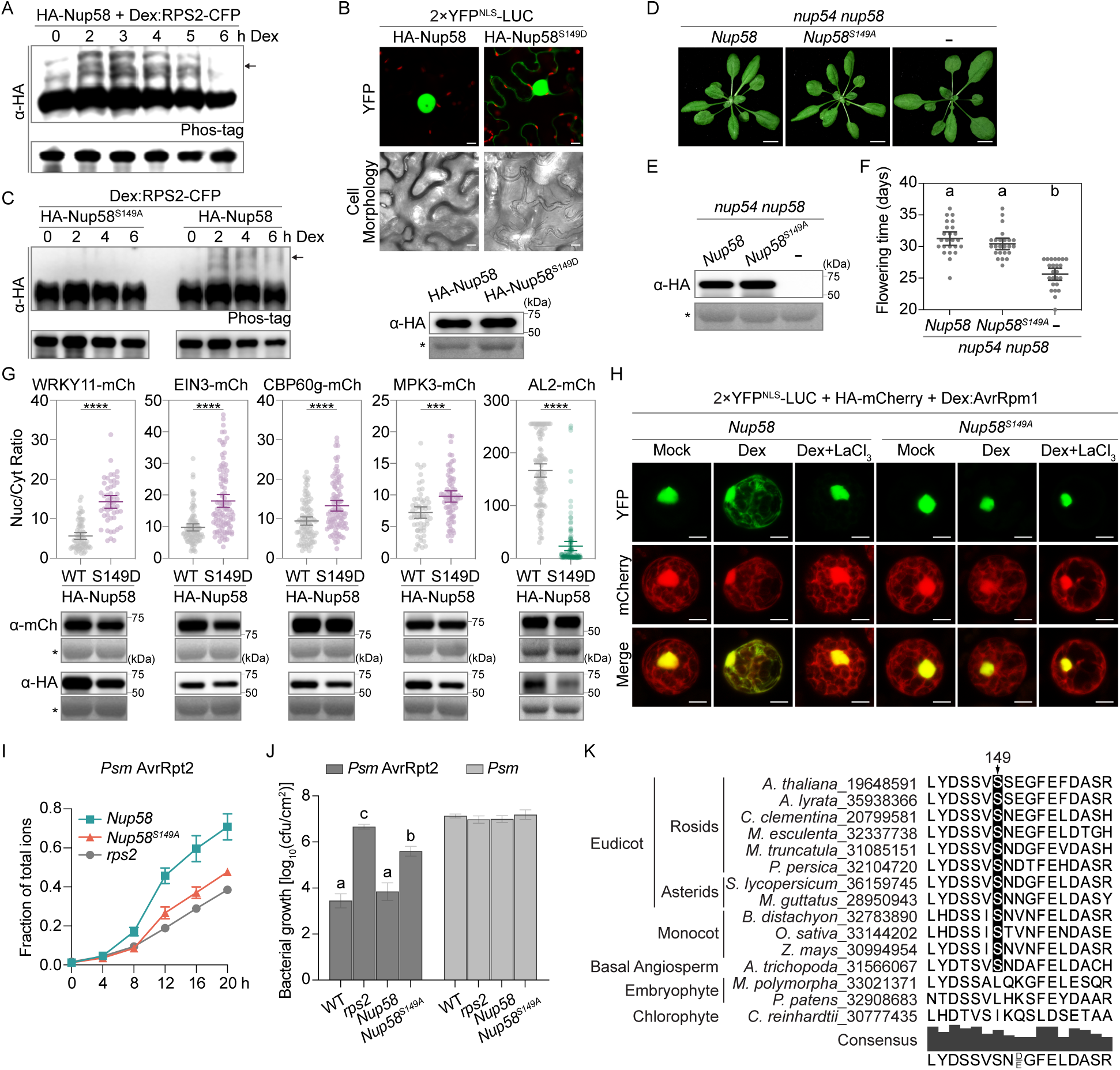
Phosphorylation of Nup58 at Ser-149 is required for the selective nuclear import of defense proteins and ETI. (A) ETI-triggered phosphorylation of Nup58. Dex-inducible RPS2 and HA-fused Nup58 were co-expressed in *N. benthamiana* leaves for 36 h and sampled after 25 μM Dex treatment at the indicated time points. The HA-Nup58 mobility shift (arrow) was detected on a Phos-tag gel using an anti-HA antibody, while immunoblotting from a normal gel (bottom) served as the loading control. (B) Cytosolic retention of the generic cargo 2×YFP^NLS^-LUC and PCD induced upon expression of the phospho-mimetic Nup58^S149D^. HA-fused Nup58 or Nup58^S149D^ was co-expressed with 2×YFP^NLS^-LUC in *N. benthamiana* leaves. YFP and cell morphology images were taken at 36 h and 60 h post-inoculation, respectively, whereas the protein levels (bottom) were detected at 36 h post-inoculation using Western blotting with an anti-HA antibody. Scale bar = 10 μm. (C) Requirement of Ser-149 in Nup58 for ETI-mediated phosphorylation. Dex-inducible RPS2 and HA-fused Nup58 or Nup58^S149A^ were co-expressed in *N. benthamiana* leaves for 36 h and ETI-mediated phosphorylation of Nup58 was examined as in (A). (D and E) The effect of the phospho-dead Nup58^S149A^ mutation on plant growth. Four-week-old transgenic *Arabidopsis* plants with either *Nup58* or *Nup58^S149A^* transformed into the *nup54 nup58* background were imaged (D) and protein levels were detected through Western blotting using an anti-HA antibody (E). Scale bar = 1 cm. (F) The effect of the phospho-dead Nup58^S149A^ mutation on the early-flowering phenotype of *nup54 nup58*. Scatter plot for days to flower, with each dot representing an individual plant and the line indicating the mean ± 95% CI. Statistical significance was assessed using one-way ANOVA followed by Tukey’s multiple comparisons test. Different lowercase letters indicate significant differences (p < 0.05). (G) Enhanced nuclear import of defense proteins triggered by the expression of phospho-mimetic Nup58^S149D^. HA-fused Nup58 (WT) or Nup58^S149D^ (S149D) was co-expressed with defense proteins or the negative control AL2 protein fused with mCherry (mCh) in *N. benthamiana* leaves for 36 h, followed by confocal imaging. The fluorescence intensity ratio between nucleus (Nuc) and cytosol (Cyt) (Nuc/Cyt ratio) was quantified (top) and the protein levels were detected through Western blotting (bottom) using an anti-mCherry or an anti-HA antibody. Each dot in the scatter plots represents a single cell, and the line represents mean ± 95% CI. WT and S149D were compared using Student’s *t*-test. ***p<0.001, ****p<0.0001. (H) Requirements of Ca^2+^ influx and Ser-149 phosphorylation in Nup58 for ETI-induced cytosolic retention of the generic cargo 2×YFP^NLS^-LUC. Plasmids carrying Dex-inducible AvrRpm1 (*Dex:AvrRpm1*), *35S:2×YFP^NLS^-LUC*, and *35S:HA-mCherry* were co-transfected into protoplasts derived from either *Nup58* or *Nup58^S149A^* transgenic plants. Confocal images were taken 2 h after water (Mock), 25 μM Dex, or 25 μM Dex plus 1 mM LaCl_3_ treatment and representative cells are shown. Scale bar = 10 μm. (I and J) The effect of the phospho-dead Nup58^S149A^ mutation on ETI-induced PCD and bacterial resistance. Four-week-old transgenic plants with either *Nup58* or *Nup58^S149A^* transformed into the *nup54 nup58* background were challenged with *Psm* AvrRpt2 at OD_600nm_ = 0.02, followed by conductivity measurements of ion leakage normalized to total ion conductivity (I). Plants were also challenged with *Psm* or *Psm* AvrRpt2 at OD_600nm_ = 0.002 and bacterial growth was measured 3 days post-inoculation (J). Data are presented as mean ± SEM (n = 3 in I and n = 8 in J). Different lowercase letters indicate significant differences based on one-way ANOVA followed by Tukey’s multiple comparisons test (p < 0.05). (K) Sequence alignment of Nup58 orthologs at Ser-149 within angiosperm plants. *, non-specific band stained by Ponceau S in protein gels.

To further examine the effect of Ser-149 phosphorylation on Nup58 activity, we transformed WT HA-Nup58 and the HA-Nup58^S149A^ variant into the *Arabidopsis nup54 nup58* double mutant background. We observed normal leaf morphology for both transgenic lines at similar protein levels (Figures 4D and 4E) and found that WT HA-Nup58 as well as the HA-Nup58^S149A^ variant could fully complement the early flowering phenotype of the *nup54 nup58* double mutant (Figure 4F), indicating that phosphorylation at Ser-149 is not required for Nup58’s function in development or flowering. In contrast, compared to WT HA-Nup58, the phospho-mimic HA-Nup58^S149D^ variant enhanced nuclear distribution of defense-related cargos, while constraining general nuclear transport, when co-expressed in *N. benthamiana* leaves (Figures 4B and 4G). Moreover, experiments using protoplasts derived from WT *Nup58* and phospho-dead *Nup58^S149A^* transgenic plants showed that treating WT protoplasts with the calcium inhibitor LaCl_3_ or using *Nup58^S149A^* protoplasts blocked effector AvrRpm1-triggered cytosolic retention of 2×YFP^NLS^-LUC observed in WT (Figure 4H). This demonstrated that phosphorylation of Nup58 at Ser-149 is required for reprogramming the NPC cargo transport during ETI in a Ca^2+^-dependent manner. Furthermore, we found that *Nup58^S149A^* transgenic plants exhibited compromised PCD induced by *Pseudomonas syringae* pv*. maculicola* ES4326 carrying the effector AvrRpt2 (*Psm* AvrRpt2) compared to the WT *Nup58* transgenic plants (Figure 4I). Consistently, higher levels of *Psm* AvrRpt2 growth were detected in the *Nup58^S149A^* transgenic plants (Figure 4J). However, when the isogenic *Psm* bacterial strain without the AvrRpt2 effector was inoculated, no detectable difference in *Psm* growth was observed between the WT *Nup58* and the *Nup58^S149A^* transgenic plants, indicating that the deficiency observed in *Nup58^S149A^* is specific to ETI. Taken together, these findings suggest that reprogramming of NPC transport activity through phosphorylation of Nup58 at Ser-149 represents a signaling event downstream of NLR activation, rather than a consequence of pathogen proliferation. The conservation of Nup58 Ser-149 in the angiosperm species (Figure 4K) provides support for the evolutionary importance of this regulatory step in plant defense. With these lines of evidence, we established a connection from ETI-induced Nup58 phosphorylation to selective nuclear import of defense proteins that ultimately leads to PCD and resistance to infection.

### CKL3-mediated phosphorylation of Nup58 is required for ETI

We next investigated all candidate kinases in the Nup58-proxiome identified through GO term MF analysis (Figure 5A), as TurboID proximity labeling can capture transient protein-protein interactions, including kinases and substrates.^36^ We selected CASEIN KINASE 1-LIKE 3 and 4 (CKL3/4) as the most likely kinases responsible for phosphorylating Nup58 at Ser-149, as they were captured by Tb-Nup58 specifically following ETI induction and are known to recognize a consensus sequence matching the region surrounding Ser-149 of Nup58.^53,54^ As Ser/Thr protein kinases, CASEIN KINASE 1 (CK1) in humans and CKLs in plants belong to an evolutionarily conserved protein kinase family in eukaryotes.^53–56^ Among the *Arabidopsis* CKL family’s 13 members, CKL3 and CKL4 have been reported to phosphorylate CRYPTOCHROME 2 (CRY2) in regulating blue light signaling.^55^ To confirm our hypothesis that CKL3 and/or CKL4 is the kinase that phosphorylates Nup58 in the NPC, we performed bimolecular fluorescence complementation (BiFC) assay by fusing CKL3 or CKL4 with the C-terminal half of YFP (cYFP), and Nup58 with the N-terminal half of YFP (nYFP). When co-expressed with Dex-inducible RPS2 in *N. benthamiana* leaves, we found that CKL3 showed a dramatically enhanced association with Nup58 upon ETI induction. In contrast, CKL4, the closest homolog of CKL3,^55^ did not show ETI-dependent increase in association with Nup58 (Figure 5B). Consistently, we observed cytosolic retention of the 2×YFP^NLS^-LUC generic reporter when co-expressed in *N. benthamiana* with the CKL3-mCherry fusion, but not with the CKL4-mCherry fusion (Figure 5C). This suggests that the ETI-triggered increase in association with Nup58, and the resulting change in nuclear transport, is largely specific to CKL3. Using pull-down and phosphorylation assays, we found that *in vitro*-expressed CKL3 could directly interact with both WT Nup58 and the Nup58^S149A^ mutant (Figure 5D), indicating that the phosphorylation status of Ser-149 does not affect Nup58 interaction with CKL3. However, the CKL3-mediated phosphorylation was diminished in the Nup58^S149A^ mutant (Figure 5E), supporting our hypothesis that CKL3 is the kinase that directly modifies Nup58 at Ser-149.

**Figure 5.**
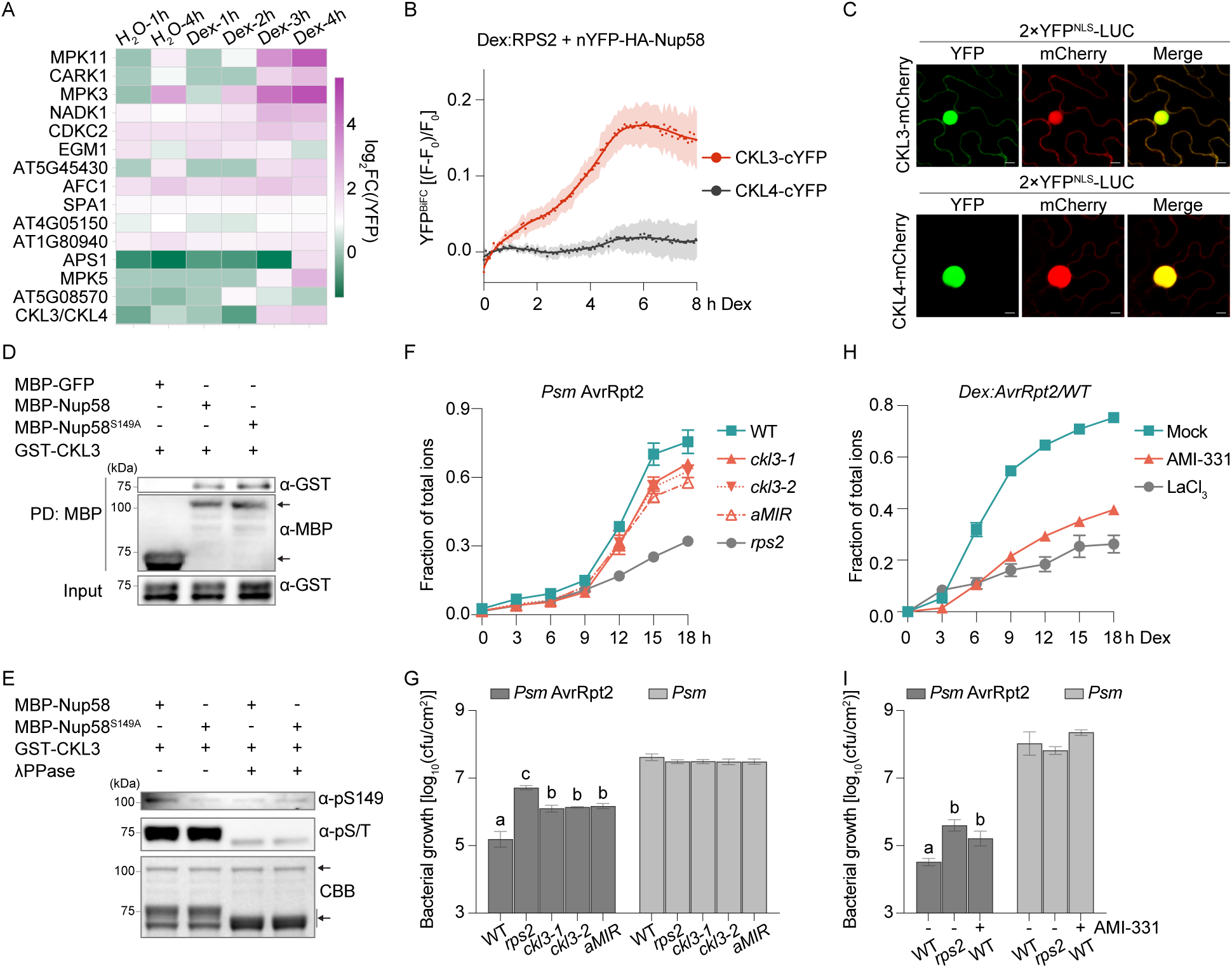
CKL3-mediated phosphorylation of Nup58 is required for ETI. (A) Heatmap depicting changes in the capture of all kinases in the Nup58-proxiome. ETI-regulated enrichment of kinases was quantified as log_2_fold change for each Tb-Nup58 sample relative to the YFP control (log_2_FC(/YFP)). (B) Bimolecular fluorescence complementation (BiFC) assay on CKL3 and CKL4 kinases for association with Nup58 during ETI. Dex-inducible RPS2 was co-expressed with Nup58 fused with the N-terminal half of YFP (nYFP-HA-Nup58) and either kinase fused with the C-terminal half of YFP (cYFP) in *N. benthamiana* leaves for 36 h. Upon Dex induction (25 μM), time-lapse recording of YFP^BiFC^ was performed on plate reader. Relative intensity was calculated based on (F-F_0_)/F_0_. (C) The effect of CKL3 or CKL4 overexpression on the cytosolic retention of the 2×YFP^NLS^-LUC reporter. CKL3 or CKL4 fused with mCherry was co-expressed with 2×YFP^NLS^-LUC in *N. benthamiana* leaves for 36 h, followed by confocal imaging with representative cells shown. Scale bar = 10 μm. (D) Pull-down assay using *in vitro*-expressed CKL3 and Nup58. MBP-Nup58, MBP-Nup58^S149A^ (top arrow), and MBP-GFP (bottom arrow) were used to pull down GST-CKL3. (E) *In vitro* phosphorylation of Nup58 by CKL3. Phosphorylation of Nup58 was detected using an antibody specific for phosphorylated Ser-149 (α-pS149), and autophosphorylation of CKL3 was detected using an anti-pS/T antibody (α-pS/T). The MBP-fused Nup58 (top arrow) and GST-CKL3 (bottom arrow) were stained by Coomassie Brilliant Blue G-250 (CBB). (F and G) The effect of *CKL3* mutation or an artificial microRNA-mediated knockdown of *CKL3/4* in a *ckl4* knockout background on ETI-associated PCD and bacterial resistance. Four-week-old plants were challenged with *Psm* AvrRpt2 at OD_600nm_ = 0.02, followed by conductivity measurements of ion leakage normalized to total ion conductivity (F). Plants were also challenged with *Psm* or *Psm* AvrRpt2 at OD_600nm_ = 0.002, followed by bacterial growth measurements 3 days post-inoculation (G). (H) The effect of the CKL-specific inhibitor, AMI-331, or Ca^2+^-channel inhibitor, LaCl_3_, on ETI-induced PCD. Four-week-old *Dex:AvrRpt2/WT* plants were pre-treated with water (Mock), 1 μM AMI-331, or 1 mM LaCl_3_ for 12 h, followed by treatment with 25 μM Dex and then conductivity measurements of ion leakage normalized to total ion conductivity at the indicated time points. (I) The effect of AMI-331 on *Psm* AvrRpt2-induced bacterial resistance. Four-week-old plants were pre-treated with and without 1 μM AMI-331 for 12 h and then challenged with *Psm* or *Psm* AvrRpt2 at OD_600nm_ = 0.002, followed by bacterial growth measurements 3 days post-inoculation. Data are presented as mean ± SEM (n = 3 in A, F, and H; n = 5 in B; and n = 8 in G and I). Different lowercase letters indicate significant differences as determined by one-way ANOVA followed by Tukey’s multiple comparisons test (p < 0.05).

To further establish the involvement of CKL3 in the ETI signaling pathway, we inoculated *Psm* AvrRpt2 into the CKL3-knockout mutants *ckl3-1* and *ckl3-2* as well as a *CKL3/4*-knockdown line expressing an artificial microRNA in the *ckl4* knockout mutant background (*aMIR*), and found that both ETI-associated PCD and bacterial resistance were partially compromised in all these *ckl* mutants (Figures 5F and 5G). Similar to the *Nup58^S149A^* plants (Figure 4J), basal resistance to *Psm* was not affected in the *ckl3* mutants (Figure 5G), indicating the specificity of CKL3 in regulating ETI. Since the *Arabidopsis* genome contains 13 *CKL*s, to overcome possible functional redundancy, we applied AMI-331, a CKL-specific inhibitor,^57^ to WT *Arabidopsis* plants and observed diminished PCD triggered by both Dex-induced AvrRpt2 and AvrRpm1, similar to the effect of the Ca^2+^ inhibitor LaCl_3_ treatment (Figures 5H and S5). In accordance with the compromised ETI/PCD, the AMI-331 treatment also resulted in more *Psm* AvrRpt2 growth (Figure 5I), supporting the requirement of CKLs for ETI-mediated resistance.

### CKL3 induces changes in the nuclear pore to reprogram transcription downstream of Ca^2+^ influx during ETI

To further pinpoint the step in the ETI signaling pathway regulated by CKL3, we examined several ETI-associated events in the presence of AMI-331. First, we found that treating *Arabidopsis* plants with AMI-331 did not negatively impact the Ca^2+^ spikes triggered by the Dex-induced expression of either AvrRpt2 or AvrRpm1, monitored by the cytoplasm-localized GCaMP3 Ca^2+^ indicator (Figures 6A and S6A). Similarly, AMI-331 treatment did not inhibit ETI-induced phosphorylation of MPK3/6 upon Dex-induced effector expression (Figures S6B and S6C). These results indicate that CKLs function either downstream of NLR-triggered Ca^2+^ influx and MPK3/6 activation or independent of these signaling events, consistent with our hypothesis that CKLs are involved in regulating Nup58 activity at the NPC. Indeed, AMI-331 treatment dampened the RPS2-triggered Nup58 phosphorylation in *N. benthamiana*, especially at 4 h after Dex-induction, as observed in the Phos-tag gel (Figure 6B). Moreover, RPS2-induced cytosolic retention of the 2×YFP^NLS^-LUC reporter was also impaired upon AMI-331 treatment, similar to the LaCl_3_ treatment which blocks Ca^2+^ influx (Figure 6C). Overall, our results suggest that CKLs (e.g., CKL3) might function as a downstream transducer of the Ca^2+^ signal generated upon ETI induction to phosphorylate Nup58 and alter NPC selectivity.

**Figure 6.**
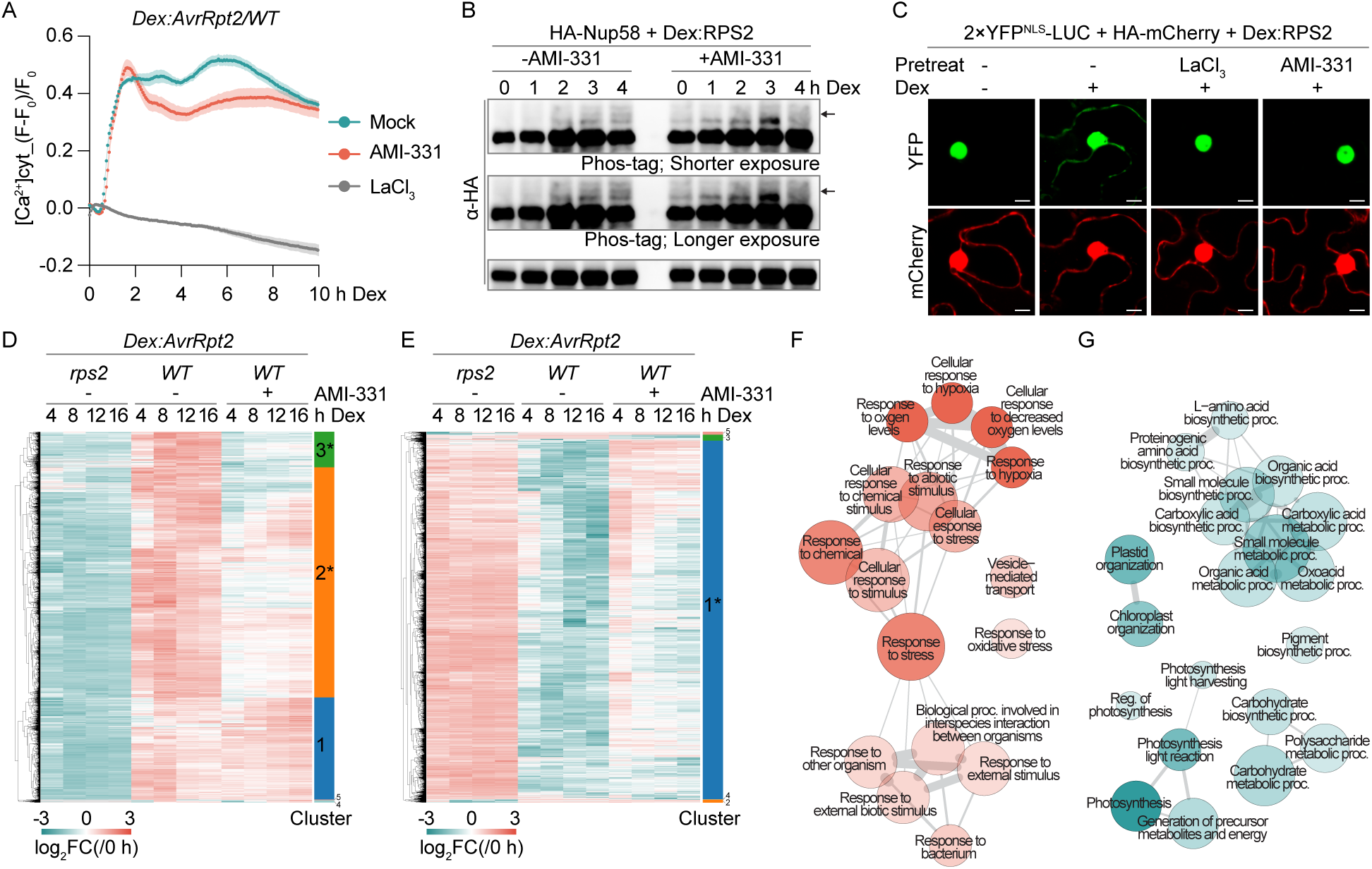
CKL3 induces changes in the nuclear pore to reprogram transcription downstream of Ca^2+^ influx during ETI. (A) The effect of AMI-331 on AvrRpt2-induced Ca^2+^ spike. Four-week-old transgenic plants expressing the cytosolic GCaMP3 Ca^2+^ indicator [Ca^2+^]cyt in the *Dex:AvrRpt2/WT* background were pre-treated with water (Mock), 1 μM AMI-331, or 1 mM LaCl_3_ for 12 h, followed by time-lapse recording of GFP fluorescence after treatment with 25 μM Dex. Cytosolic Ca^2+^ concentration dynamics were calculated based on (F-F_0_)/F_0_. Data are presented as mean ± SEM (n = 10). (B) The effect of AMI-331 on ETI-induced Nup58 phosphorylation. The Dex-inducible RPS2 and HA-fused Nup58 were co-expressed in *N. benthamiana* leaves for 24 h. Leaf discs were then pre-treated with and without 1 μM AMI-331 for 12 h. Samples were collected at the indicated time points after 25 μM Dex treatment. HA-Nup58 mobility shift (arrow) was detected on a Phos-tag gel using an anti-HA antibody, while immunoblotting from a normal gel (bottom) was used as the loading control. (C) The effect of AMI-331 on ETI-induced cytosolic retention of the generic cargo 2×YFP^NLS^-LUC. *N. benthamiana* leaves were co-inoculated with *Agrobacteria* carrying *Dex:RPS2*, *35S:2×YFP^NLS^-LUC*, and *35S:HA-mCherry* for 24 h. Leaf discs were then pre-treated with and without 1 mM LaCl_3_ or 1 μM AMI-331 for 12 h. Confocal images were taken after 3 h, with and without 25 μM Dex treatment with representative cells shown. Scale bar = 10 μm. (D) The effect of AMI-331 on ETI-mediated gene induction. Clustered heatmap of ETI up-regulated Differentially Expressed Genes (ETI-UP DEGs) was generated using the log_2_fold change of each gene normalized to that at time zero (log_2_FC(/0 h)). * indicates clusters exhibiting reduced ETI induction following AMI-331 pre-treatment. (E) The effect of AMI-331 on ETI-mediated gene repression. Clustered heatmap of ETI down-regulated DEGs (ETI-DN DEGs) was generated using log_2_FC(/0 h). * indicates the cluster exhibiting diminished ETI repression after AMI-331 pre-treatment. (F and G) GO networks of AMI-331-impacted ETI-UP DEGs (F) and ETI-DN DEGs (G), constructed using ShinyGO 0.85.1 from DEGs in clusters 2 and 3 (D) and cluster 1 (E), respectively, and visualized in R.

Within the Nup58-proxiome, “regulation of transcription” ranks among the most enriched GO terms (Figures 2F and S2E). It led us to propose a simple model: CKL3 phosphorylates Nup58, which enhances selective cargo transport through the nuclear pore. This, in turn, activates ETI-associated transcriptional changes that result in PCD and pathogen resistance. To test this hypothesis, we performed a quantitative 3’ mRNA sequencing (QuantSeq) in a Dex-induced ETI time course (Figure S6D) using transgenic *Arabidopsis* plants expressing *Dex:AvrRpt2* in the WT background with and without AMI-331 pre-treatment, and in the *rps2* mutant background without AMI-331 pre-treatment. To identify ETI-regulated transcripts, we first applied Weighted Gene Co-expression Network Analysis (WGCNA) on gene expression data comparing WT and *rps2* without AMI-331 pre-treatment, resulting in 12 co-expression modules with distinct expression patterns (Figure S6E). Using the conductivity dynamics observed in Figure S6D, which reflect the progression of ETI-associated PCD, as the phenotypic trait, we then examined how each module’s eigengene correlated with the conductivity measurements as well as the candidate hub genes in each module which show high correlation with both the module eigengene and the phenotypic trait. We found two modules, Module 1 and 12, exhibiting both the highest absolute Pearson Correlation Coefficients (|*r*| > 0.5) and the largest proportions of candidate hub genes within the modules (HubRatio > 0.5) (Figure S6F). Based on their eigengene profiles, we designated the candidate hub genes from Module 1 as ETI up-regulated DEGs (ETI-UP DEGs, Table S4) and the candidate hub genes from Module 12 as ETI down-regulated DEGs (ETI-DN DEGs, Table S5). To examine the impact of AMI-331 on ETI-mediated transcription in WT plants, we generated clustered heatmaps using the log_2_fold change of each gene normalized to that at time zero (log_2_FC(/0 h)) within ETI-UP DEGs (Figure 6D) and ETI-DN DEGs (Figure 6E). We observed that 2139 of 2984 genes (71.7%) from ETI-UP DEGs (Clusters 2 and 3) showed reduced ETI induction following AMI-331 pre-treatment (Figures 6D and S6G, Table S4), while 4886 of 5051 genes (96.7%) from ETI-DN DEGs (Cluster 1) displayed less ETI-triggered repression with AMI-331 pre-treatment (Figures 6E and S6H, Table S5). More interestingly, the AMI-331-impacted ETI-UP DEGs were highly enriched in stress response-related terms (Figure 6F), whereas the AMI-331-impacted ETI-DN DEGs were enriched in photosynthesis, chloroplast organization, and amino acid biosynthesis-related terms (Figure 6G), consistent with the GO terms identified in previous ETI transcriptomic analyses.^15,58,59^ The large proportion of ETI-regulated DEGs being affected by AMI-331 provides strong support for our hypothesis that ETI-induced transcriptional reprogramming is largely mediated by CKL3. Together with the compromised PCD and bacterial resistance observed in the *Nup58^S149A^* transgenic plants (Figures 4I and 4J), in the *ckl3* mutants (Figures 5F and 5G), and after AMI-331 treatment (Figures 5H, 5I and S5), these results provide genetic as well as mechanistic evidence that CKL3-mediated phosphorylation of Nup58 alters NPC permeability, and that the resulting transcriptional reprogramming is essential for ETI-associated PCD and pathogen resistance.

### Ca^2+^-binding promotes CKL3-Nup58 association for enhanced defense protein nuclear import and ETI execution

Since inhibition of CKL activity by AMI-331 does not impair NLR-mediated Ca^2+^ influx (Figures 6A and S6A), CKL3-mediated phosphorylation of Nup58 likely occurs downstream of Ca^2+^ influx and may be directly regulated by it. Indeed, LaCl_3_ treatment could block ETI-enhanced association between CKL3 and Nup58 without impacting their protein abundance (Figures 7A and 7B). To explore the potential role of Ca^2+^ in promoting CKL3-Nup58 interaction, we used AlphaFold3 to model Ca^2+^-binding to CKL3, which identified Asp-149 in CKL3 as a potential Ca^2+^-binding residue through formation of two hydrogen bonds; substitution with alanine (D149A) abolished this interaction (Figure 7C). Consistently, *in vitro*-expressed CKL3 was shown to bind to Ca^2+^ with a *K*_d_ of 12.2 ± 5.9 μM, while the CKL3^D149A^ variant failed to bind Ca^2+^, in the Microscale Thermophoresis assay (Figure 7D). Moreover, ETI-enhanced association with Nup58 was also abolished in the CKL3^D149A^ variant in transient expression analysis using *N. benthamiana* (Figures 7E and 7F), indicating that the NLR-induced Ca^2+^ influx is required for CKL3 to interact with its substrate, Nup58.

**Figure 7.**
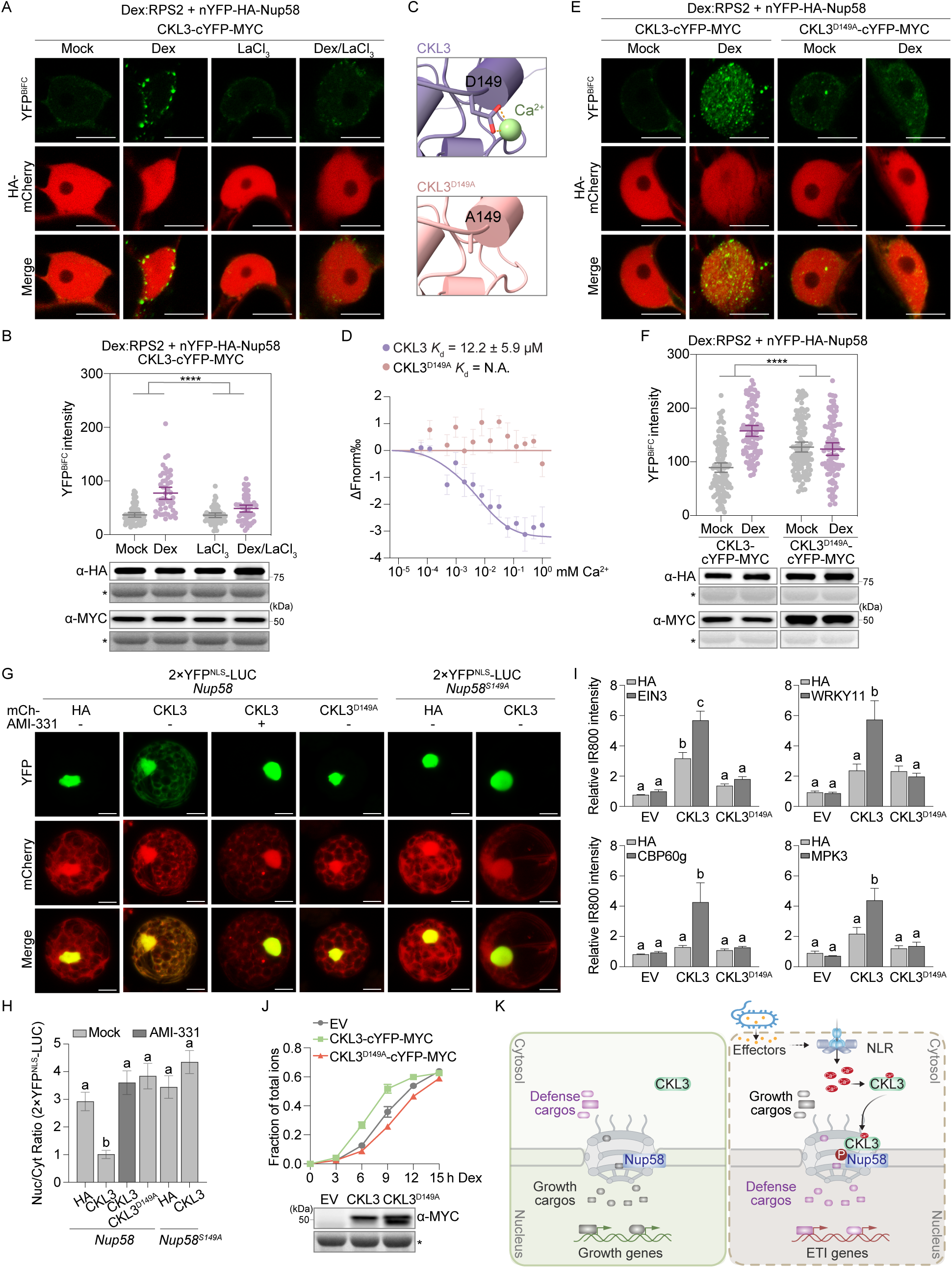
Ca^2+^-binding promotes CKL3-Nup58 association for enhanced defense protein nuclear import and ETI execution. (A and B) ETI-enhanced, Ca^2+^-dependent association between CKL3 and Nup58. Dex-inducible RPS2 and HA-mCherry were co-expressed with nYFP-HA-Nup58 and CKL3-cYFP-MYC in *N. benthamiana* leaves for 36 h. YFP^BiFC^ images were taken 3 h after 25 μM Dex treatment with and without 1 mM LaCl_3_ (A), and YFP^BiFC^ intensities were quantified (B, top) with protein level detected through Western blotting using an anti-HA or an anti-MYC antibody (B, bottom). Each dot in the scatter plot represents a single cell, and the line represents mean ± 95% CI. Two-way ANOVA was performed, ****p<0.0001. The asterisk (*) in protein gels marks non-specific band stained by Ponceau S. (C) AlphaFold3 prediction of Asp-149 in CKL3 as a Ca^2+^-binding residue. (D) Microscale thermophoresis analysis of Ca^2+^ binding to *in vitro*-expressed WT CKL3 and CKL3^D149A^. Each binding assay was independently performed eight times, and ΔFnorm‰ was used for data analysis across replicates. Solid curves indicate fitted data, and dots represent mean ± SEM (n = 8). *K*_d_, dissociation constant; N.A., not available. (E and F) The effect of the CKL3^D149A^ mutation on ETI-enhanced association between Nup58 and CKL3. Dex-inducible RPS2 and HA-mCherry were co-expressed with nYFP-HA-Nup58, CKL3-cYFP-MYC or CKL3^D149A^-cYFP-MYC in *N. benthamiana* leaves for 36 h. YFP^BiFC^ images were taken 3 h post water (Mock) or 25 μM Dex treatment (E) and YFP^BiFC^ intensities were quantified (F, top) with protein level detected through Western blotting using an anti-HA or an anti-MYC antibody (F, bottom). Each dot in the scatter plot represents a single cell, and the line represents mean ± 95% CI. Two-way ANOVA was performed, ****p<0.0001. The asterisk (*) in protein gels marks non-specific band stained by Ponceau S. (G and H) The effect of Nup58 phosphorylation and Ca^2+^-binding on CKL3-triggered cytosolic retention of 2×YFP^NLS^-LUC. Plasmids carrying *35S:2×YFP^NLS^-LUC* and mCherry fusions (*35S:HA-mCherry*, *35S:CKL3-mCherry* or *35S:CKL3^D149A^-mCherry*) were co-transfected into protoplasts derived from either *Nup58* or *Nup58^S149A^* transgenic plants and treated with and without 1 μM AMI-331. Confocal images were taken 12 h after incubation (G, scale bar = 10 um) and fluorescence intensity ratio between nucleus (Nuc) and cytosol (Cyt) (Nuc/Cyt ratio) of 2×YFP^NLS^-LUC was quantified (H). Data are presented as mean ± SEM (n ≥ 20). Different lowercase letters indicate significant differences using one-way ANOVA followed by Tukey’s multiple comparisons test (p < 0.05). (I) The effect of co-expressing cargos from the UP group on CKL3-mediated, Ca^2+^-dependent cell death. HA-fused mCherry (HA), or a cargo from the UP group (EIN3, WRKY11, CPB60g or MPK3) was co-expressed with CKL3, CKL3^D149A^, or the empty vector (EV) in *N. benthamiana* leaves and cell death-associated autofluorescence was measured using Infrared Fluorescence imaging (IR800) 7 days post-inoculation with intensity qualified in ImageJ. Data are presented as mean ± SEM. Different lowercase letters indicate significant differences based on one-way ANOVA followed by Tukey’s multiple comparisons test (p < 0.05). (J) The effect of expressing CKL3 on ETI-induced cell death. Dex-inducible RPS2 was co-expressed with either CKL3-cYFP-MYC (CKL3), CKL3^D149A^-cYFP-MYC (CKL3^D149A^) or the empty vector (EV) in *N. benthamiana* leaves for 36 h, followed by conductivity measurements of ion leakage normalized to total ion conductivity at the indicated time points after treatment with 25 μM Dex (top). The protein levels were detected prior to Dex treatment through Western blotting using an anti-MYC antibody (bottom). Data are presented as mean ± SEM (n = 3). The asterisk (*) in protein gels marks non-specific band stained by Ponceau S. (K) Model illustrating the signaling pathway from NLR resistosome-triggered Ca^2+^ influx to Ca^2+^-dependent CKL3 interaction with and phosphorylation of Nup58, resulting in preferential nuclear import of defense cargos, transcriptional reprogramming, and execution of ETI.

To examine the functional consequence of the Ca^2+^-mediated CKL3 interaction with Nup58, we overexpressed mCherry-fused (mCh) CKL3 or CKL3^D149A^ together with the 2×YFP^NLS^-LUC reporter in protoplasts derived from either *Nup58* or *Nup58^S149A^ Arabidopsis* transgenic plants (Figure 4D). We observed that overexpression of CKL3 led to cytosolic retention of the 2×YFP^NLS^-LUC reporter in *Nup58* protoplasts, but not in *Nup58^S149A^* protoplasts (Figures 7G and 7H). Additionally, this effect on the 2×YFP^NLS^-LUC transport in *Nup58* protoplasts was abolished following AMI-331 treatment and was absent when the Ca^2+^-binding-deficient CKL3^D149A^ was overexpressed. Similar results were obtained in *N. benthamiana* through transient overexpression of CKL3 versus CKL3^D149A^ (Figure S7A). Together, these findings establish a signaling cascade, in which NLR-mediated Ca^2+^ influx enhances CKL3’s interaction with Nup58, leading to Nup58 phosphorylation at Ser-149 and thereby altering NPC selectivity.

Finally, to link the Ca^2+^-mediated CKL3 function in facilitating preferential nuclear import of defense proteins to the execution of ETI, we generated transgenic plants with Dex-inducible CKL3 or CKL3^D149A^. We found that repeated Dex treatment to overexpress CKL3 from the seedling stage markedly impaired plant growth and development, likely due to spontaneous cell death, whereas plants overexpressing the Ca^2+^-binding-deficient CKL3^D149A^ grew similarly to WT (Figure S7B). To determine whether CKL3-induced PCD phenotype is due to its activity in altering the transport activity of the nuclear pore in response to ETI, we co-expressed UP-group cargos from the Nup58-proxiome (Figure 3, Table S1), together with either WT CKL3 or the Ca^2+^-binding-deficient CKL3^D149A^ variant in *N. benthamiana*. We found that co-expression of these UP-group cargos with WT CKL3 further enhanced CKL3-mediated PCD, whereas co-expression with the CKL3^D149A^ variant did not (Figure 7I), even at comparable protein levels (Figure S7C). Importantly, the enhanced PCD observed for the UP-group cargos was not detected for the DN-group cargo, AL2 (Figures S7D and S7E), consistent with our hypothesis that overexpression of CKL3 mimics the ETI-induced state and the resulting selective nuclear import of defense-related cargos causes the observed PCD. Moreover, CKL3 overexpression in *N. benthamiana* accelerated PCD triggered by Dex-induced RPS2 expression, an effect not observed with the vector control or the CKL3^D149A^ variant (Figure 7J), further supporting a role for Ca^2+^-dependent CKL3 activity in promoting ETI-associated PCD. Notably, the conservation of Asp-149 in CKL3 across *Arabidopsis* CKL members, angiosperm CKL3 orthologs, and human CK1δ/ε (Figures S7F and S7G) suggests that CKLs and CK1s may have broader Ca^2+^-dependent regulatory functions beyond mediating nuclear transport via Nup58 phosphorylation during the ETI response in plants.

## DISCUSSION

Recent advances in understanding how NLR resistosomes assemble, generate nucleotide-derived secondary messengers, and trigger sustained Ca^2+^ influx have highlighted a significant gap in our knowledge of their downstream signaling mechanisms. While PCD has been used as a major output of NLR-mediated ETI, there have been evidence indicating that execution of ETI relies on a combination of multiple responses in the cytoplasm and the nucleus before the onset of PCD and full-scale resistance.^7,9^ Following resistosome-mediated Ca^2+^ influx, ETI also involves ROS production through the activity of NADPH oxidase on the plasma membrane,^60^ activation of various intracellular kinases such as MAPKs and CDPKs,^41,61^ transcriptional reprogramming in the nucleus,^15,24,58,59^ and an increase in translation which is associated with ATP-mediated assembly of the eukaryotic translation initiation factor 2 complex.^15,16^ The involvement of multiple regulatory mechanisms makes it difficult to identify downstream signaling components using classical genetic approaches, as mutations in these components will often produce only partial or subtle ETI phenotypes. To overcome this obstacle and establish the signaling link between cytosolic Ca^2+^ influx and nuclear transcriptional reprogramming, we used the nuclear pore channel protein Nup58 as a probe to capture the ETI-induced changes in the NPC activity in an unbiased manner. Surprisingly, contrary to our previous hypothesis that the NPC is deregulated during ETI,^24^ the induced NPC instead selectively transports specific defense proteins while restricting general import mediated by the canonical NLS mechanism. We show that this ETI-mediated switch in the nuclear pore selectivity is triggered through phosphorylation of Nup58 at Ser-149 by the Ca^2+^-bound CKL3, which ultimately leads to ETI-associated transcription, PCD, and disease resistance (Figure 7K).

The NPC, as the gateway between cytoplasm and nucleoplasm, has been associated with various cellular processes. During apoptosis in animals, large cargo proteins, which are normally excluded from the nucleus, were found in the nucleus through more permeable nuclear pores.^62^ During glutamate-induced neuronal cell death, both size-exclusion and facilitated nuclear import became defective.^63^ It was also found that nuclei of old rat neurons contained deteriorated NPCs with increased nuclear permeability and accumulation of cytoplasmic tubulin.^64^ In plants, multiple Nups have been reported to impact defense responses, with distinct effects on cargo transport.

Overexpression of CPR5, a membrane-associated Nup, led to more constrained nuclear import;^24^ a loss of *Nup88* function induced nuclear protein export;^22^ and mutating the Nup107-160 complex caused reduced mRNA export.^23^ Our discovery of the Ca^2+^-CKL3-Nup58 signaling cascade demonstrates that the NPC permeability is regulated during ETI in a precisely controlled manner, not by passive breakdown of the barrier, and ETI-associated transcription and cell death are highly choreographed events at the NPC. How phosphorylation of Nup58 at Ser-149 alters the selective permeability of defense cargos, while constraining the general NLS-mediated nuclear transport remains unclear. We have ruled out cargo MW and isoelectric point, as well as transcriptional and translational regulation of cargo genes, as major driving mechanisms (Figure S3). It is possible that “calmodulin-binding”, an enriched GO term of the UP^85^ group (Figure 2F), may explain some of the cargos’ preferential transport. Notably, the IDR-containing proteins are overrepresented within the NPC-associated proteome, suggesting a phase-separated environment surrounding the NPC.^37^ Recent studies have shown that phase separation of central Nups mediates nuclear transport of the kinase MPK3 to promote resistance against *Botrytis cinerea*, whereas the pathogen effector BcSSP2 suppresses NPC phase separation to enhance virulence.^26,65^ Whether ETI-mediated changes in NPC selectivity involves phase separation warrants further investigation.

The effect of Ca^2+^ on casein kinase activity has not been well documented. Unlike the Ca^2+^-dependent kinases, *in vitro*-expressed casein kinase does not rely on Ca^2+^ to phosphorylate its substrates (Figure 5E).^54–56^ In many cases, scaffold proteins, like the FAM83 family proteins in mammals, are needed for casein kinases to become proximal to their substrates *in vivo*.^56^ In this study, we showed that *in vitro*-expressed CKL3 exhibited affinity to Ca^2+^ through Asp-149 (Figure 7D), which is conserved not only among all the CKL members in *Arabidopsis* and within angiosperm plants, but also in human CK1δ/ε (Figures S7F and S7G). During ETI, CKL3 association with Ca^2+^ enhances its interaction with Nup58 (Figures 7A and 7B), which leads to the phosphorylation of Nup58. This ETI-enhanced association is abolished when the Ca^2+^-binding site is mutated in CKL3^D149A^ (Figures 7E and 7F). Whether this Ca^2+^-enhanced interaction is substrate-specific or general needs further investigation. The Ca^2+^-dependent regulation being a broadly conserved feature across other CKLs and CK1s is an exciting possibility.

In this study, we demonstrate that precise nucleocytoplasmic transport is paramount for successful defense responses. The enhanced nuclear import of defense-related TFs and transcriptional regulators (Figure 3) provides a direct mechanistic link to ETI-associated transcriptional reprogramming. Multiple defense-related proteins captured in our Nup58-proxiome likely act collectively to reprogram transcription. Disrupting this process at the level of Nup58 phosphorylation, via *Nup58^S149A^*, or CKL3 inhibition, provides an opportunity to address the importance of transcriptional reprogramming in executing PCD and pathogen resistance during ETI (Figures 4H, 6B-6G, 7G and 7H). The impaired ETI-associated PCD phenotype observed in the *Nup58^S149A^* transgenic plants, the *ckl* mutants, and upon treatment with the CKL-specific inhibitor AMI-331 (Figures 4I, 4J, 5F-5I and S5) supports the current view that full execution of ETI requires both cytoplasmic and nuclear functions. Our findings reveal a signaling cascade connecting resistosome-induced cytosolic Ca^2+^ influx to nuclear transcriptional reprogramming and identify CKL3, Nup58, and its associated cargos as new entry points for understanding ETI execution in plants.

## ACKNOWLEDGEMENTS

We thank Dr. Zhi-yong Wang for sharing the *YFP-YFP-TurboID* line; Dr. Hongwei Xue for sharing the *amiR-CKL3/4.ckl4* lines (*aMIR*) and GST-CKL3/pGEX-4T-1 construct; Dr. Daoxin Xie for sharing the pETL7-NUP58 and pETL7-EGFP constructs; Dr. David Mackey for sharing the *Dex:AvrRpm1/rpm1 rps2* line; Dr. Pei Zhou and Karly Forker for assisting the setup of MST experiments. We thank Dr. Jordan Powers for helping discussion on the project and other Dong lab members for critical reading of the manuscript. This work was supported by grants from the National Institutes of Health (R35-GM118036-06); National Science Foundation (IOS-2041378), and the Howard Hughes Medical Institute to X.D. This work was also supported by grants from the National Institutes of Health (S10OD030441) to S.-L.X. and by the Carnegie Endowment Fund to the Carnegie Mass Spectrometry Facility.

## AUTHOR CONTRIBUTIONS

X.Z. generated most of the data presented in this study. A.V.R. and S.-L.X. performed the LC-MS/MS for the TurboID samples. S.K. assisted with QuantSeq library construction. S.K. and Y.C.X. assisted with QuantSeq data analysis. Y.Z.X. performed the phylogenetic analysis. X.Z. and X.D. wrote the manuscript with contributions from all coauthors.

## DECLARATION OF INTERESTS

X.D is a co-founder of Upstream Biotechnology Inc. and a member of its scientific advisory board, as well as a scientific advisory board member of Inari Agriculture Inc and Aferna Bio.

## RESOURCE AVAILABILITY

### Lead contact

Further information and requests for reagents may be directed to and will be fulfilled by the corresponding author Xinnian Dong.

### Materials availability

All unique materials and constructs in this study are available from the lead contact upon completion of Material Transfer Agreement.

### Data and code availability

- All data reported in this paper will be shared by the lead contact upon request.
- The LC-MS/MS proteomics data have been deposited into the ProteomeXchange Consortium via PRIDE^66^ partner repository with the dataset identifier PXD077161.
- QuantSeq sequencing data are available through NCBI under accession number PRJNA1450108.

## STAR★METHODS

## EXPERIMENTAL MODEL AND SUBJECT DETAILS

### Plants and growth conditions

All *Arabidopsis thaliana* plants used in this study were in the Columbia-0 (WT) ecotype background, grown at 22 ℃ in a 12 h /12 h light/dark photoperiod with 60% relative humidity for 3-4 weeks before experiments. *Nicotiana benthamiana* plants were grown under the same conditions for 5-6 weeks before experiments. *Arabidopsis* lines *Dex:AvrRpt2/WT*,^67^ *Dex:AvrRpt2/rps2*,^24^ *35S:YFP-YFP-TurboID*,^36^ *nup54 nup58*,^68^ *rps2*,^69^ and *amiR-CKL3/4.ckl4* (*aMIR*)^55^ were previously described. T-DNA insertion mutants *ckl3-1* (Salk_016571C) and *ckl3-2* (Salk_110494C) were obtained from the Arabidopsis Biological Resource Center (ABRC). The *35S:TurboID-3*×*HA-Nup58* transgenic lines were generated in the *Dex:AvrRpt2/WT* background; *35S:HA-Nup58* and *35S:HA-Nup58^S149A^* transgenic lines in the *nup54 nup58* background; and *Dex:CKL3-mCherry* and *Dex:CKL3^D149A^-mCherry* transgenic lines in the WT background. The calcium imaging lines were generated by transforming *35S:GCaMP3^NES^* into the *Dex:AvrRpt2/WT* background; or co-transformed with *Dex:AvrRpm1* into the WT background. The *Dex:AvrRpm1/WT* lines were generated by crossing *Dex:AvrRpm1/rpm1 rps2*^70^ and WT, followed by PCR-based genotyping of the F2 population.

## METHOD DETAILS

### Plasmid construction

Multisite Gateway Assembly was performed as described^24^ with adjustments: the full-length coding sequence (CDS) of TurboID, sYFP2, and the N-terminal YFP (nYFP) were cloned into the pBSDONR p1-p4 vector; 3×HA, sYFP2, and sYFP2^NLS^ were cloned into the pBSDONR p4r-p3r vector; and the full-length cDNAs of *Nup58* and *luciferase (LUC)* were cloned into the pBSDONR p3-p2 vector, using the BP reaction (BP clonase, Thermo Scientific) to generate the entry clones. For constructs with constitutive expression under the 35S promoter: (1) the LUC/p3-p2 entry clone was paired with sYFP2/p1-p4 plus either sYFP2/p4r-p3r or sYFP2^NLS^/p4r-p3r entry clones to generate the fusion constructs 2×YFP-LUC and 2×YFP^NLS^-LUC, respectively, in the pEG100 destination vector via LR reaction (LR clonase II plus, Thermo Scientific); and (2) the Nup58/p3-p2 entry clone was paired with 3×HA/p4r-3r plus TurboID/p1-p4, 3×HA/p1-p4, or nYFP/p1-p4 entry clones to generate fusion constructs of TurboID-3×HA-Nup58 (Tb-Nup58), 3×HA-3×HA-Nup58 (HA-Nup58) and nYFP-3×HA-Nup58 (nYFP-HA-Nup58), respectively, in the pEG100 destination vector via LR reaction. For constructs driven by the Dex-inducible promoter: (1) full-length cDNAs of *RPS2*, *RPM1*, *NRG1.D485V,* and *AvrRpm1* were cloned into pBSDONR p1-p4 through BP reaction, and paired with sCFP3A/p4r-p2, 3×HA/p4r-p2, and 5×MYC/p4r-p2 entry clones to generate fusion constructs of RPS2-CFP, RPS2-HA or AvrRpm1-HA, and RPM1-MYC or NRG1.DV-MYC, respectively, in the pBAV154 destination vector by LR reaction and (2) the cDNA of *RPS2* was cloned into the pBSDONR p4r-p2 vector through BP reaction and paired with the sCFP3A/p1-p4 entry clone to generate CFP-RPS2 fusion in the pBAV154 destination vector by LR reaction.

HiFi DNA Assembly: the full-length cDNAs of *AL2*, *LNK1*, *RPN10*, *WRKY11*, *EIN3*, *CAMBP25*, *CBP60g*, *SARD1*, *MPK3*, *CKL3*, and *CKL4* were PCR-amplified and combined with mCherry or cYFP-MYC coding sequence to be inserted into the Xho1/Spe1-digested pEG100 or pBAV154 destination vectors for constitutive or Dex-inducible expression, using the NEBuilder HiFi DNA Assembly Master Mixture (New England BioLabs Inc.). Full-length CDS of GCaMP3^NES^ was assembled with the Xho1/Xba1-digested pEG100 vector to generate *35S:GCaMP3^NES^* and transform into the *Dex:AvrRpt2/WT* background; or was assembled with the BamH1/Pst1-digested pCambia1300 vector, with the resulting *35S:GCaMP3^NLS^:E9 Ter* cassette from the pCambia1300 vector further PCR-amplified and inserted into the Spe1-digested AvrRpm1-HA/pBAV154 fragment to generate *35S:GCaMP3^NES^-Dex:AvrRpm1* and co-transformed into WT background. All point mutations were introduced by PCR-mediated mutagenesis and confirmed through sequencing.

### Transient expression in *N. benthamiana* and analyses

*Agrobacterium*-mediated transient expression in *N. benthamiana* was performed as described^71^ with minor adjustments. In brief, *Agrobacterium* suspensions were adjusted to OD_600nm_ = 0.2 for the NLR-expressing strains and OD_600nm_ = 0.6 for all other strains; then combined in equal volumes before infiltrating into fully expanded *N. benthamiana* leaves. For the cargo localization assay and the bimolecular fluorescence complementation (BiFC) assay, leaf discs were subjected to the indicated treatments 36 h post-infiltration, followed by confocal imaging using a Carl Zeiss 880 Airyscan inverted confocal laser scanning microscope with a 488 nm laser plus 516-522 nm emission filter for the YFP signal and a 561 nm laser plus 592-629 nm emission filter for the mCherry signal. Cell morphology was imaged 60 h post-infiltration using the confocal microscope under white field. Dead cell-associated autofluorescence was imaged 1 week post-infiltration on an Azure biosystems using a 784 nm laser and 832 nm emission filter for IR800. All confocal images were acquired under non-saturating conditions and analyzed by ImageJ. To calculate the cargo distribution, straight lines covering either the nucleus or the cytoplasm were drawn within the same cell to measure the fluorescence intensity in the two compartments using the Region of Interest Manager in ImageJ (as shown in Figure 3) to calculate the Nuc/Cyt ratio. For time-lapse BiFC, leaf discs were subjected to Dex solution; YFP^BIFC^ signals were collected on plate reader (PerkinElmer, VICTOR NIVO^TM^) with a 480/30 nm excitation filter plus a 530/30 nm emission filter. The average fluorescence intensity from the first five timepoints was taken as base line (F_0_), and the relative intensity at each time point, (F-F_0_)/F_0_, was calculated.

### Measurement of ETI responses

To quantitatively measure cell death in *N. benthamiana* 36 h post-inoculation, leaf discs from infiltrated regions were soaked in a 25 μM dexamethasone (Dex, Sigma-Aldrich) solution, and the conductivity resulting from ion leakage was measured using an Orion Star series meter (Thermo Scientific). To measure cell death in *Dex:AvrRpt2/WT, Dex:AvrRpt2/rps2,* or *Dex:AvrRpm1/WT* transgenic *Arabidopsis,* leaf discs from 4-week-old plants were soaked for 12 h in water (Mock), in 1 μM AMI-331 (TCI Chemicals), or in 1 mM lanthanum chloride (LaCl_3_, Sigma-Aldrich) before the addition of 25 μM Dex. Conductivity was then measured. For ETI-associated cell death triggered by bacteria, *Pseudomonas syringae* pv. *maculicola* ES4326 carrying AvrRpt2 (*Psm* AvrRpt2)^72^ were grown on King’s B medium for 2 days and then infiltrated into leaves of 4-week-old plants (OD_600nm_ = 0.02 in 10 mM MgCl_2_). Leaf discs were collected and soaked in water for conductivity measurements. For bacterial growth measurements, 4-week-old plants were infiltrated with bacterial suspensions of *Psm* or *Psm* AvrRpt2 (OD_600nm_ = 0.002 in 10 mM MgCl_2_), and colony-forming units (cfu) were counted 3 days later.

### TurboID proximity labeling and purification of biotinylated proteins

Four-week-old transgenic plants expressing TurboID-3×HA-Nup58 (Tb-Nup58) in the Dex-inducible AvrRpt2 (*Dex:AvrRpt2/WT*) background were sprayed with a 25 μM Dex solution for 1, 2, 3 and 4 h to induce ETI or with H_2_O for 1 and 4 h as the background to identify ETI-specific targets. To first remove random background signals from Dex-induced and uninduced samples, 4-week-old transgenic plants constitutively expressing 2×YFP-TurboID (YFP), known to be ubiquitously distributed in both cytoplasm and nucleus,^36^ were also sprayed with a 25 μM Dex solution for 4 h. To biotinylate TurboID-proximal proteins, a 50 μM biotin solution was sprayed 3 h prior to sample collection, which was carried out at the same time for all samples with three replicates (∼ 6 g/replicate) for each sample and time point. Affinity purification of biotinylated proteins was performed as previously described^73^ with modifications. Briefly, finely ground plant tissues were fully dissolved in 9 mL of extraction buffer [50 mM Tris-HCl pH 7.5, 150 mM NaCl, 1% SDS, 1% Triton X-100, 0.5% Na-deoxycholate, 1 mM EGTA, 1 mM DTT, 1× protease inhibitor cocktail (cOmplete Tablets Mini EDTA-free *EASYpack*, Roche) and 1× phosphatase inhibitor cocktail (PhosSTOP *EASYpack*, Roche)]. Protein extracts were collected from the clear supernatant obtained after filtering, sonication, and centrifugation. To remove excess free biotin, 3 mL of the protein extract was loaded onto a PD-10 desalting column (Cytiva) which was equilibrated with 25 mL of ice-cold equilibration buffer (50 mM Tris-HCl pH 7.5, 150 mM NaCl, 0.1% SDS, 1% Triton X-100, 0.5% Na-deoxycholate, 1 mM EGTA, 1 mM DTT), and eluted with 3.5 mL of elution buffer (50 mM Tris-HCl pH 7.5, 150 mM NaCl, 0.01% SDS, 1% Triton X-100, 0.5% Na-deoxycholate, 1 mM EGTA, 1 mM DTT, 1× protease inhibitor cocktail and 1× phosphatase inhibitor cocktail). Dynabeads MyOne Streptavidin C1 (Invitrogen) slurry (200 μL) pre-washed with the elution buffer were added to 10.5 mL of final elute. After incubating in the cold room for 16 h, beads were washed as described^73^ and resuspended in 1 mL of the equilibration buffer supplemented with 1× protease inhibitor cocktail and 1× phosphatase inhibitor cocktail, and then subjected to Mass Spec analysis.

### LC-MS/MS and data analysis

Trypsin digestion and LC-MS/MS were carried out as described previously.^71^ Protein identification and label-free quantification (LFQ) were performed in MaxQuant 2.0.3.1 using default settings. Filtering was then performed with Perseus 1.6.15.0 to remove proteins marked as “only identified by site”, “reverse”, and “potential contaminant”. LFQ values were log_2_-transformed and those proteins with at least two valid values in at least one sample were kept for further analyses. Missing values were imputed using the function of “Replace missing values from normal distribution”. A two-sample *t*-test was performed between the Nup58 samples (H_2_O- and Dex-treated) and the YFP control to identify significant Nup58 partners, defined by -log_10_(p-value) > 1 and log_2_FC(/YFP) > 1 (Figure S2D, Table S1) and partners from all six treatments/timepoints were then combined to generate the Nup58-proxiome. Gene Ontology (GO) term enrichment was performed using Protein Analysis Through Evolutionary Relationships (PANTHER)^74^ for Figure S2E and Database for Annotation, Visualization and Integrated Discovery (DAVID, https://davidbioinformatics.nih.gov/tools.jsp) for Figure 2. Principal Component Analysis (PCA) and Clustered heatmap were performed using SRplot (https://www.bioinformatics.com.cn/en) and plots were generated using Prism 10 and SRplot, respectively.

### Phos-tag gel assay and Western blotting

Six *N. benthamiana* leaf discs were collected after the indicated treatment and ground into fine powder, followed by suspension in 100-150 μL of SDS buffer (200 mM Tris-HCl pH 6.8, 6% SDS, 30% glycerol, 15% bromophenol blue and 15% β-mercaptoethanol), incubation on ice for 30 min, heating at 95 ℃ for 10 min, and centrifugation at 4 ℃ for 10 min. For Phos-tag gel assay, the supernatant was loaded onto 8% SDS-polyacrylamide gels containing 40 μM MnCl_2_ and 50 μM Phos-tag Acrylamide (FUJIFILM Wako). After electrophoresis, proteins were wet-transferred to the nitrocellulose (NC) membrane using the transfer buffer containing 0.1% SDS, followed by immunoblotting with an anti-HA antibody (Cell Signaling). For normal gel assay, the supernatant was loaded on 4-12% NuPAGE Bis-Tris Protein Gels (Invitrogen) and proteins were dry-transferred to the NC membrane, followed by immunoblotting with an anti-HA antibody (Cell Signaling), an anti-MYC antibody (Cell Signaling), an anti-mCherry antibody (Chromotek), or an anti-pMPK3/6 antibody (Cell Signaling).

### Phylogenetic analysis

Ortholog groups of AtNup58 and AtCKL3 were identified as previously described,^75^ and presented with species name abbreviation_PACID. Protein sequences from each species were retrieved from Phytozome 13.^76^ Protein sequences for *Arabidopsis* CKLs and human CK1δ/ε were retrieved from National Center for Biotechnology Information (NCBI) and presented with species name abbreviation_GI number. Multiple sequence alignment was performed using MAFFT implemented in the EMBL-EBI Job Dispatcher sequence analysis tool^77^ with the following parameters: BLOSUM62 substitution matrix, gap open penalty of 1.53, and gap extension penalty of 0.123. The results were visualized using AlignmentViewer (http://alignmentviewer.org) with the indicated peptide presented.

### Antibody preparation

Peptide of LYDSSV-pS-SEGFEF-Cys was used to generate the phospho-specific antibodies against Nup58^S149^ (anti-pS149) by PACIFIC Immunology.

### *In vitro* pull-down assay

GST-CKL3 expression in the BL21 culture was induced by 1 mM isopropylthio-β-galactoside (IPTG) when OD_600nm_ reached 0.8 and pellets were collected after shaking at 28 ℃ for 3 h. MBP-fused proteins (MBP-GFP, MBP-Nup58 and MBP-Nup58^S149A^) were also expressed in BL21 and induced by 0.5 mM IPTG when OD_600nm_ reached 0.8. Pellets were then collected after shaking at 16 ℃ for 16 h. GST-CKL3 and MBP-fused proteins were purified using Pierce^TM^ Glutathione Agarose (Thermo Scientific) and Amylose Resin (New England BioLabs Inc.), respectively, based on manufacturer’s instructions with minor adjustment. Pellets for GST-CKL3 and MBP-fused proteins were extracted using Low Salt Buffer (20 mM Tris-HCl pH 8.0, 150 mM NaCl, 10% glycerol, 1 mM DTT, 1 mM EDTA, 0.2% Triton X-100, 0.5 mM PMSF, 1× protease inhibitor cocktail and 1× phosphatase inhibitor cocktail) and High Salt Buffer (40 mM Tris-HCl pH 8.0, 500 mM NaCl, 10% glycerol, 1 mM DTT, 0.5 mM PMSF, 1× protease inhibitor cocktail and 1× phosphatase inhibitor cocktail), respectively, followed by sonication. The clear supernatant after centrifugation was exposed to the indicated resins and incubated in the cold room for 12 h. Proteins were eluted by incubating with Low Salt Buffer + 10 mM GSH for GST-CKL3 and High Salt Buffer + 10 mM Maltose for MBP-fused proteins, after washing with Low Salt Buffer or High Salt Buffer for 5 times. The products were concentrated using Amicon Ultra-4 spin columns.

Purified MBP-fused proteins (10 μg) were incubated with 15 μL of Amylose Resin in 800 μL High Salt Buffer at 4 ℃ for 12 h. After washing the beads with High Salt Buffer for 3 times, 50 μg of purified GST-CKL3 was added and incubated further at 4 ℃ for 12 h. Beads were washed for 5 times with High Salt Buffer and finally resuspended in the SDS buffer followed by Western blotting with an anti-MBP antibody (Invitrogen) and an anti-GST antibody (Cytiva).

### *In vitro* phosphorylation assay

Purified GST-CKL3 (50 μg) was incubated with 10 μg of purified MBP-Nup58 or MBP-Nup58^S149A^ in 40 μL of kinase buffer (50 mM Tris-HCl pH 8.0, 10 mM MgCl_2_, 1 mM DTT, 20 mM ATP) at 25 ℃ for 3 h. Then the 40 μL kinase reaction was split into two tubes with one for λ

PPase digestion [4 μL kinase buffer, 3 μL MnCl_2_, 3 μL NEBuffer Pack, and 1 μL λPPase (Lambda Protein Phosphatase, New England BioLabs Inc.)] and the other as the control (4 μL kinase buffer, 3 μL MnCl_2_, and 3 μL NEBuffer Pack). Both tubes were incubated at 37 ℃ for 1 h and the reactions were terminated by adding 30 μL of the SDS buffer. Samples were then heated at 95 ℃ for 10 min followed by Western blotting using the Nup58^S149^ phospho-specific antibodies (anti-pS149, PACIFIC Immunology) and an anti-pS/T antibody (Cell Signaling). Coomassie Brilliant Blue G-250 (CBB) was used to stain MBP-Nup58 and MBP-Nup58^S149A^ at ∼ 100 kDa as well as GST-CKL3 at ∼ 75 kDa.

### Microscale Thermophoresis (MST) assay

MST for quantifying Ca^2+^-binding affinity was described previously^78^ with modifications. GST-CKL3^D149A^ was similarly expressed and purified *in vitro* as GST-CKL3. Purified GST-CKL3 and GST-CKL3^D149A^ (10 μM) in 100 μL were labeled by Monolith Protein Labeling Kit RED-NHS 2^nd^ Generation (NanoTemper Technologies Inc.) for 30 min in the dark at room temperature following manufacturer’s instructions. The labeled protein was diluted to 50 nM using the MST buffer (50 mM Tris-HCl pH 8.0, 150 mM NaCl, 0.05% Tween-20 and 0.1% Pluronic F-127). Meanwhile, a 16-point 1:1 serial dilution of CaCl_2_ was prepared in the MST buffer, with the resulting CaCl_2_ gradient ranging from 1 mM to 30.5 nM. 10 μL of 50 nM labeled proteins were mixed with 10 μL of CaCl_2_ solutions and incubated in dark at room temperature for 30 min, then loaded into Monolith Premium Capillaries (NanoTemper Technologies Inc.). Binding was measured using Monolith X (NanoTemper Technologies Inc.) at 25 ℃, 100% Excitation and High IR Laser Power. Curve fitting and dissociation constant (*K*_d_) calculation were performed using the MO.Control v2.7.1 software (NanoTemper Technologies Inc.). Each binding assay was repeated eight times independently and ΔFnorm‰ was used to incorporate multiple replicates. The graph was generated using Prism 10.

### QuantSeq and data analysis

Leaf discs from 4-week-old *Dex:AvrRpt2/WT* and *Dex:AvrRpt2/rps2* transgenic plants were soaked with and without 1 μM AMI-331 for 12 h prior to Dex treatment (25 μM). Samples were collected at the indicated hours after Dex addition. RNA extraction and QuantSeq library prep were performed as previously described.^71^ Weighted Gene Co-expression Network Analysis (WGCNA)^79^ was performed using *WT* and *rps2* samples in R (with the softpower being 8), and 12 modules in total were obtained. Using the conductivity dynamics as the phenotypic trait, two parameters were assessed to identity the ETI-regulated transcripts: (1) the correlation between each module’s eigengene and the phenotypic trait (absolute Pearson Correlation Coefficients > 0.5); and (2) the fraction of candidate hub genes within each module (HubRatio > 0.5). The candidate hub genes were selected when their absolute correlation coefficients with the module’s eigengene were > 0.8 and their absolute correlation coefficients with the phenotypic trait were > 0.5. Altogether, candidate hub genes from Module 1 and Module 12 were designated as ETI up-regulated DEGs (ETI-UP DEGs) and ETI down-regulated DEGs (ETI DN-DEGs), respectively. Clustered heatmaps were generated in R, based on the log_2_fold change of each gene normalized to that at time zero (log_2_FC(/0 h)). GO network was built in ShinyGO 0.85.1 and graphed in R, with the darker color indicating more significantly enriched gene sets, the bigger nodes indicating larger gene sets, and thicker edges indicating more overlapped genes (only connected if 20% or more genes were shared).

### Calcium imaging and quantification

Leaf discs from 4-week-old *GCaMP3^NES^*/*Dex:AvrRpt2* and *GCaMP3^NES^*/*Dex:AvrRpm1* transgenic plants were soaked in water (Mock), 1 μM AMI-331 or 1 mM LaCl_3_ for 12 h prior to Dex treatment (25 μM). GCaMP3^NES^ signals were imaged in time-lapse with 3.667 min interval, using the fluorescence stereo microscope Leica M205FA with a 450-490 nm excitation filter plus a 500-550 nm emission filter. Fluorescence intensity (F) at each time point was measured using ImageJ. The average fluorescence intensity from first five timepoints was taken as base line (F_0_) and the relative intensity at each time point (F-F_0_)/F_0_ was calculated.

## QUANTIFICATION AND STATISTICAL ANALYSIS

All experiments were repeated at least two times with similar results. Data plotting and statistical tests were performed using Prism 10, with the statistical information such as mean ± SEM (standard error), 95% CI (confidence intervals), and significance testing, specified in the figure legends.

