## Supplemental Figures for "Ca^2+^-activated CKL3 phosphorylates nucleoporin 58 to reprogram nuclear transport and execute effector-triggered immunity"

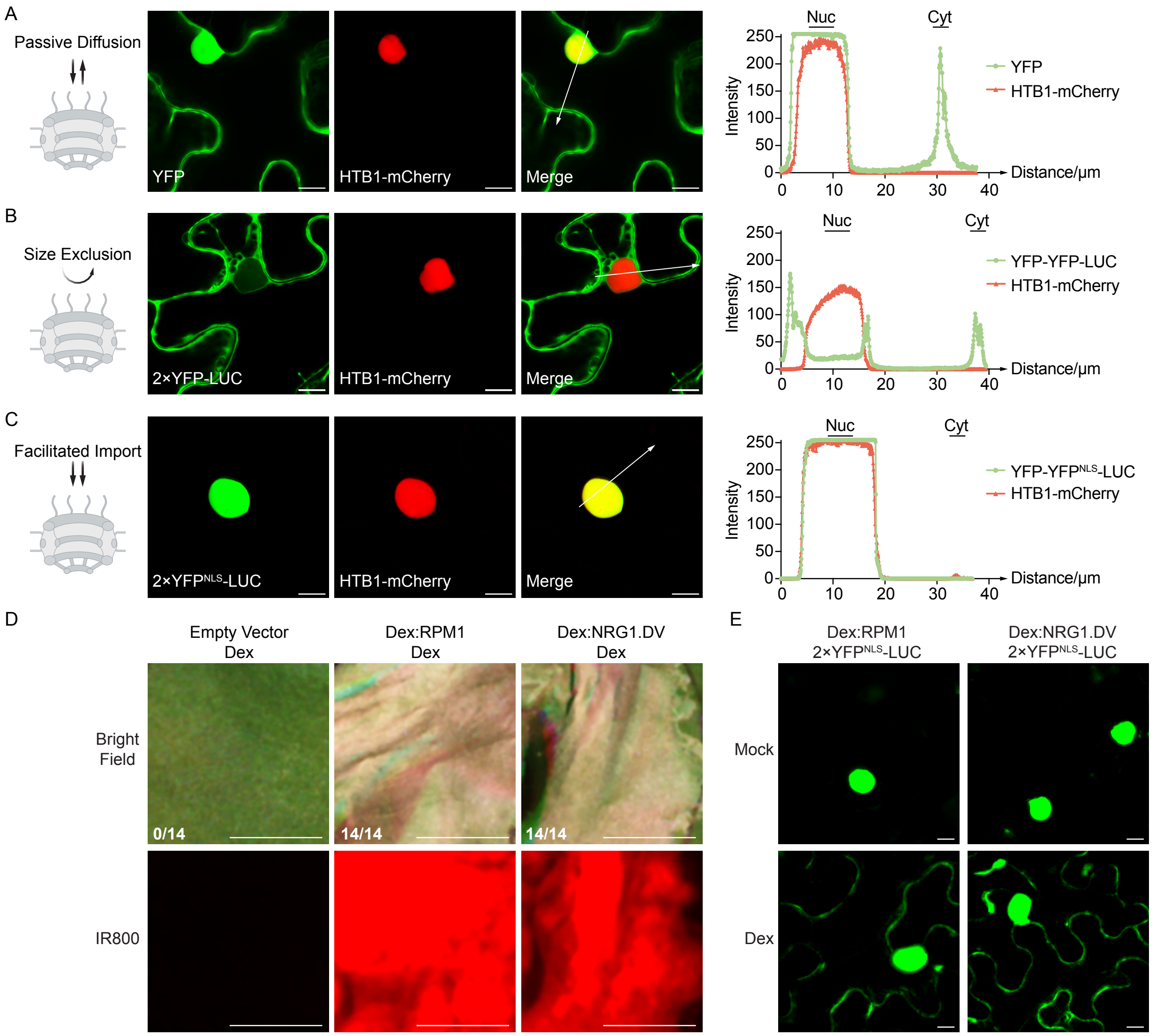


**Figure S1. NLRs induce changes in nuclear import and PCD. Related to Figure 1.**

(A-C) The design of fluorescent reporters to assess different modes of transport activities. *N. benthamiana* leaves were inoculated with *Agrobacterium* carrying *35S:YFP*, *35S:2×YFP-LUC*, or *35S:2×YFP$^{NLS}$-LUC* together with *35S:HTB1-mCherry* (HTB1, a histone marker for the nucleus) for 36 h followed by confocal imaging. To represent different interactions with the nuclear pore (left), fluorescent reporters were imaged (middle) to detect passive diffusion of YFP (27 kDa) between the cytosol and the nucleus (A); size exclusion by the nuclear pore of 2×YFP-LUC (117 kDa) with cytosolic localization (B); and facilitated nuclear import of 2×YFP$^{NLS}$-LUC with nuclear localization (C). Fluorescence intensities along the arrows covering both the nuclear and the cytosolic positions were measured and graphed (right). Scale bar = 10 μm.

(D) RPM1- or NRG1.DV-induced PCD. After inoculating *N. benthamiana* with *Agrobacterium* carrying *Dex:RPM1* or *Dex:NRG1.DV* as in (A-C), the plants were treated with 25 μM Dex for 1 day followed by counting dead versus total inoculated leaves under bright field (top) and by measuring Infrared Fluorescence (IR800) to visualize cell death-associated autofluorescence (bottom). Scale bar = 0.5 cm

(E) Cytosolic retention of the generic nuclear transport reporter 2×YFP$^{NLS}$-LUC induced by RPM1 or NRG1.DV. *N. benthamiana* leaves were inoculated with *Agrobacterium* carrying *Dex:RPM1* or *Dex:NRG1.DV* together with *35S:2×YFP$^{NLS}$-LUC* for 36 h and images were taken 3 h post water (Mock) or 25 μM Dex (Dex) treatment. Scale bar = 10 μm.

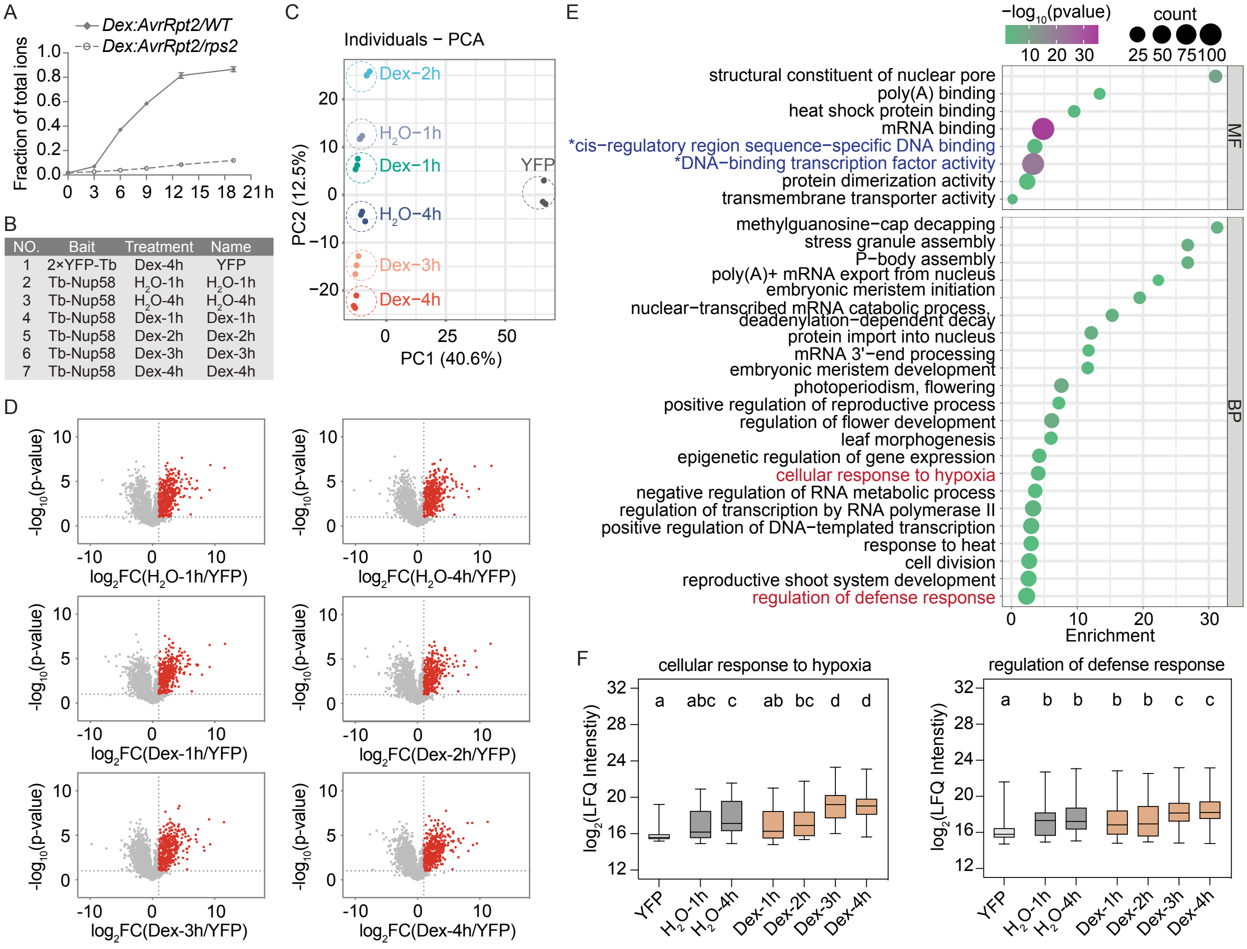


| NO. | Bait | Treatment | Name |
|---|---|---|---|
| 1 | 2×YFP-Tb | Dex-4h | YFP |
| 2 | Tb-Nup58 | $H_2O$-1h | $H_2O$-1h |
| 3 | Tb-Nup58 | $H_2O$-4h | $H_2O$-4h |
| 4 | Tb-Nup58 | Dex-1h | Dex-1h |
| 5 | Tb-Nup58 | Dex-2h | Dex-2h |
| 6 | Tb-Nup58 | Dex-3h | Dex-3h |
| 7 | Tb-Nup58 | Dex-4h | Dex-4h |

**Figure S2. Tb-Nup58 captures cargos passing through the nuclear pore during ETI. Related to Figure 2.**

(A) Time-course measurements of ETI-triggered PCD in *Arabidopsis* plants. Four-week-old transgenic plants carrying *Dex:AvrRpt2* in the wild-type (WT) and AvrRpt2-receptor mutant *rps2* background were treated with 25 μM Dex. Conductivity, used as a proxy for ion leakage from cells undergoing PCD, was recorded and normalized to total ion conductivity.

(B) Baits and treatments used in the time-course Nup58-TurboID experiment and their abbreviated names.

(C) Principal component analysis (PCA) of Nup58 and YFP samples. Colored circles denote the three replicates for each sample.

(D) Nup58-specific targets under 6 treatment conditions. Each Nup58 sample was compared to the YFP control to identify Nup58-specific targets with $log_2FC(/YFP) > 1$ and $-log_{10}(p\text{-value}) > 1$, indicated by red dots in the Volcano plots. Targets from all 6 treatments constitute the Nup58-proxiome.

(E) Gene Ontology (GO) enrichment of molecular function (MF) and biological process (BP) terms for the Nup58-proxiome, using the *Arabidopsis* proteome as the background.

(F) Box-and-whisker plots showing the two BP categories from (E, red) with increased capture by Tb-Nup58 upon ETI induction. Repeated-measures one-way ANOVA with Greenhouse-Geisser correction, followed by Tukey's multiple comparisons test, was performed for each category based on $log_2$(LFQ Intensity). Different lowercase letters indicate significant differences ($p < 0.05$).

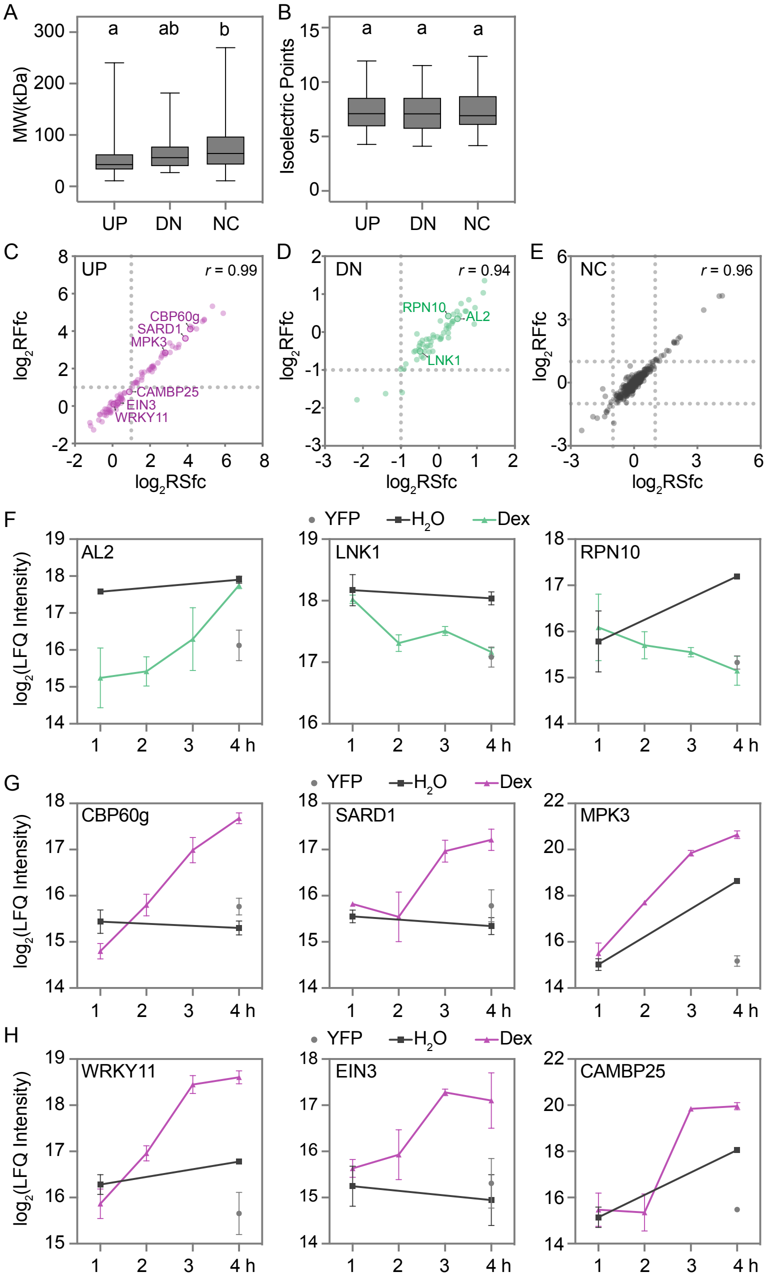


**Figure S3. ETI enhances nuclear import of defense-related proteins while constraining general cargo transport. Related to Figure 3.**

(A and B) Box-and-whisker plots of molecular weight (MW) (A) and isoelectric point (B) of proteins in the UP, Down (DN), and Non-changed (NC) groups of the Nup58-proxiome. Different lowercase letters indicate significant differences from a one-way ANOVA followed by Tukey's multiple comparisons test ($p < 0.05$).

(C-E) ETI-induced transcriptional and translational changes in the Nup58-proxiome UP (C), DN (D), and NC (E) groups, based on data from a previous RNA-seq and Ribo-seq study.[15] RSfc, fold change in RNA-seq. RFfc, fold change in Ribo-seq. *r*, Pearson correlation coefficient.

(F-H) Dynamic changes in cargos captured by Tb-Nup58 during ETI in the DN group (F), the UP group with ETI-mediated transcriptional/translational upregulation (G), and the UP group without ETI-mediated transcriptional/translational upregulation (H). Data are presented as mean ± SEM ($n = 3$).

A

SES[134]QRLDQCSR

LYDS[146]S[147]VS[149]S[150]EGFEFDASR

B

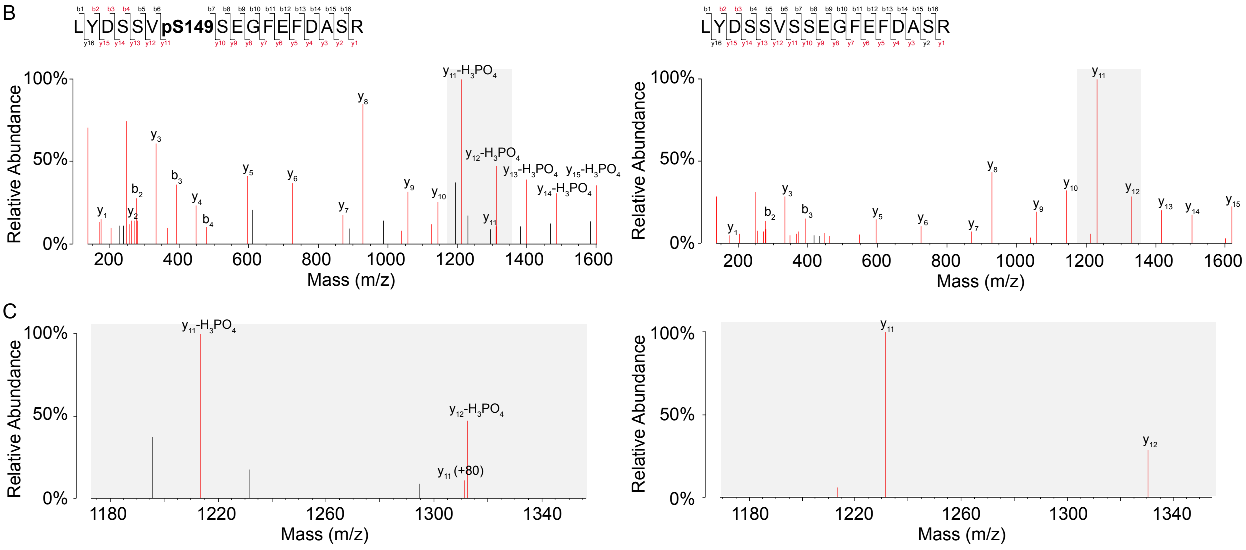


D

Main Sequence Ions (m/z)

| b | | | | y | y-$H_3PO_4$ |
|---|---|---|---|---|---|
| --- | 1 | L | 17 | --- | --- |
| 277.1547 | 2 | Y | 16 | 1862.7116 | 1764.7347 |
| 392.1816 | 3 | D | 15 | 1699.6483 | 1601.6714 |
| 479.2136 | 4 | S | 14 | 1584.6214 | 1486.6445 |
| 566.2457 | 5 | S | 13 | 1497.5893 | 1399.6124 |
| 665.3141 | 6 | V | 12 | 1410.5573 | 1312.5804 |
| 832.3124 | 7 | pS149 | 11 | 1311.4889 (+80) | 1213.5120 |
| 919.3445 | 8 | S | 10 | 1144.4905 | --- |
| 1048.3871 | 9 | E | 9 | 1057.4585 | --- |
| 1105.4085 | 10 | G | 8 | 928.4159 | --- |
| 1252.4769 | 11 | F | 7 | 871.3945 | --- |
| 1381.5195 | 12 | E | 6 | 724.3260 | --- |
| 1528.5879 | 13 | F | 5 | 595.2835 | --- |
| 1643.6149 | 14 | D | 4 | 448.2150 | --- |
| 1714.6520 | 15 | A | 3 | 333.1881 | --- |
| 1801.6840 | 16 | S | 2 | 262.1510 | --- |
| --- | 17 | R | 1 | 175.1190 | --- |

Main Sequence Ions (m/z)

| b | | | | y |
|---|---|---|---|---|
| --- | 1 | L | 17 | --- |
| 277.1547 | 2 | Y | 16 | 1782.7453 |
| 392.1816 | 3 | D | 15 | 1619.6820 |
| 479.2136 | 4 | S | 14 | 1504.6550 |
| 566.2457 | 5 | S | 13 | 1417.6230 |
| 665.3141 | 6 | V | 12 | 1330.5910 |
| 752.3461 | 7 | S | 11 | 1231.5226 |
| 839.3781 | 8 | S | 10 | 1144.4905 |
| 968.4207 | 9 | E | 9 | 1057.4585 |
| 1025.4422 | 10 | G | 8 | 928.4159 |
| 1172.5106 | 11 | F | 7 | 871.3945 |
| 1301.5532 | 12 | E | 6 | 724.3260 |
| 1448.6216 | 13 | F | 5 | 595.2835 |
| 1563.6486 | 14 | D | 4 | 448.2150 |
| 1634.6857 | 15 | A | 3 | 333.1881 |
| 1721.7177 | 16 | S | 2 | 262.1510 |
| --- | 17 | R | 1 | 175.1190 |

E

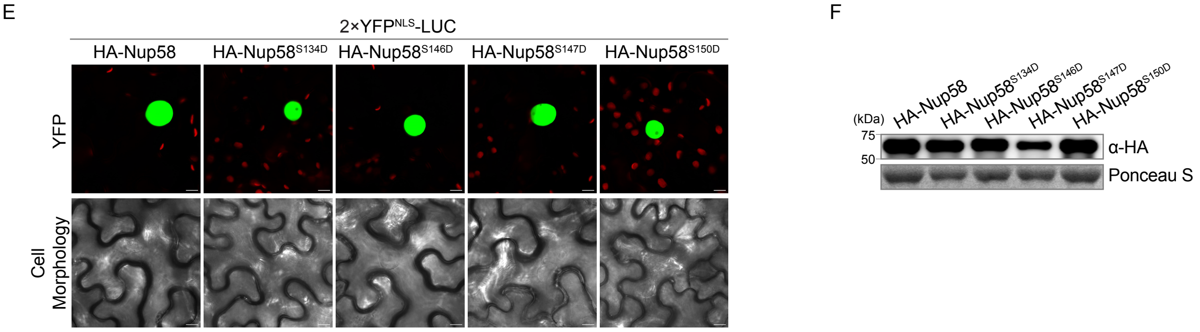


**Figure S4. Phosphorylation of Nup58 at Ser-149 is required for the selective nuclear import of defense proteins and ETI. Related to Figure 4.**

(A) Two phospho-peptides of Nup58 detected by TurboID LC-MS/MS during ETI, with putative phosphorylation sites marked in red.

(B-D) MS/MS spectra for Ser149-phosphorylated peptides (left) and non-phosphorylated peptides (right). The grey boxed regions in the full spectra (B) were enlarged in (C). Main sequence ions (m/z) were listed in (D), with the detected ions marked with red.

(E and F) Impacts of Nup58 phospho-mimic mutants on nuclear transport and PCD. HA-tagged Nup58 or a mutant variant was co-expressed with 2×YFP$^{NLS}$-LUC in *N. benthamiana* leaves for 36 h, followed by confocal imaging (E) and Western blotting with an anti-HA antibody (F). Scale bar = 10 μm.

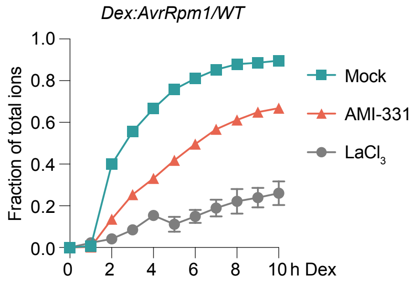


**Figure S5. CKL3 is a candidate kinase involved in ETI. Related to Figure 5.** The effect of the CKL-specific inhibitor AMI-331 or $Ca^{2+}$ inhibitor $LaCl_3$ on ETI-triggered PCD. Four-week-old *Dex:AvrRpm1/WT* plants were pre-treated with water (Mock), 1 µM AMI-331, or 1 mM $LaCl_3$ for 12 h, followed by time-course conductivity measurements of ion leakage normalized to total ion conductivity after Dex treatment (25 µM). Data are presented as mean ± SEM (n = 3).

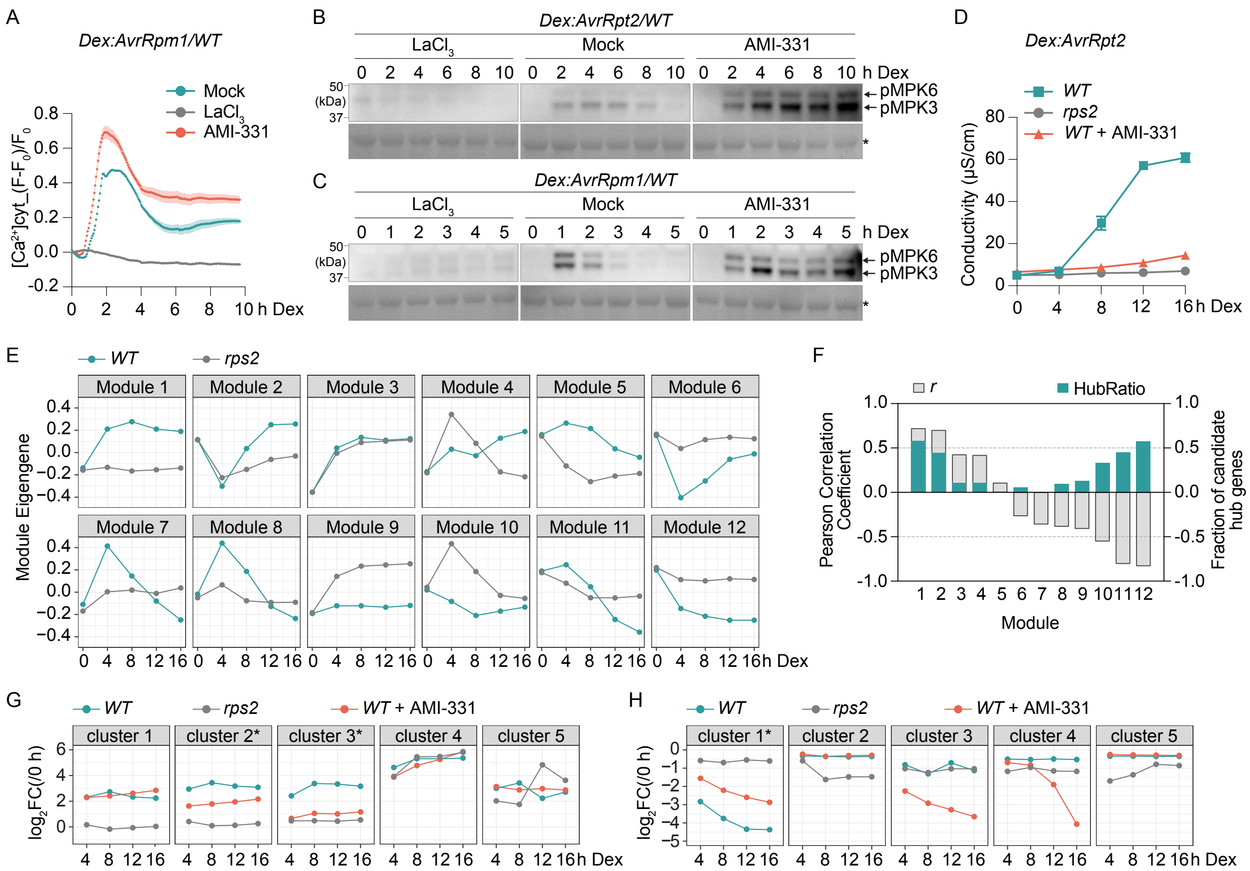


**Figure S6. CKL3 induces changes in the nuclear pore to reprogram transcription downstream of $Ca^{2+}$ influx during ETI. Related to Figure 6.**

(A) The effect of the CKL-specific inhibitor AMI-331 on the AvrRpm1-induced $Ca^{2+}$ spike. Four-week-old transgenic plants expressing the cytosolic $Ca^{2+}$ indicator GCaMP3$^{NES}$ ($[Ca^{2+}]$cyt) in the *Dex:AvrRpm1/WT* background were pre-treated with water (Mock), 1 μM AMI-331, or 1 mM $LaCl_3$ for 12 h, followed by GFP time-lapse imaging upon 25 μM Dex treatment. $Ca^{2+}$ concentration dynamics were calculated by the relative intensity based on $(F-F_0)/F_0$. Data are presented as mean ± SEM (n = 11).

(B and C) The effect of the CKL-specific inhibitor AMI-331 on MPK3/6 phosphorylation induced by AvrRpt2 (B) or AvrRpm1 (C). Four-week-old transgenic plants carrying *Dex:AvrRpt2* or

*Dex:AvrRpm1* were pre-treated with water (Mock), 1 μM AMI-331, or 1 mM $LaCl_3$ for 12 h, followed by 25 μM Dex treatment. Samples were collected at the indicated time points post Dex treatment and examined using Western blotting with an antibody for phosphorylated MPK. *, non-specific bands stained by Ponceau S.

(D) Conductivity assay performed in parallel to the QuantSeq experiment. Four-week-old transgenic plants carrying *Dex:AvrRpt2* in either WT or the *rps2* background were pre-treated with and without 1 μM AMI-331 for 12 h, followed by 25 μM Dex treatment. At indicated time points post Dex induction, samples were collected for QuantSeq and conductivity was measured in parallel. Data are presented as mean ± SEM (n = 3).

(E) Co-expression modules generated by Weighted Gene Co-expression Network Analysis (WGCNA) of QuantSeq data from WT and *rps2* samples without AMI-331 pre-treatment. Module eigengenes were plotted for each module.

(F) Pearson correlation coefficients (*r*) between module eigengenes (E) and the PCD phenotypic trait (D), and fraction of candidate hub genes (HubRatio) in each module. Modules 1 (induced) and 12 (repressed) have both high $|r|$ (> 0.5, left y-axis) and high HubRatio (> 0.5, right y-axis).

(G and H) The effect of the CKL-specific inhibitor AMI-331 on ETI-regulated transcription. The averaged $\log_2$fold change of each gene normalized to that at time zero ($\log_2$FC(/0 h)) for the five clusters from ETI-UP DEGs was plotted in (G) and that for the five clusters from ETI-DN DEGs was plotted in (H), respectively. * indicates the clusters showing compromised ETI regulation upon AMI-331 pre-treatment.

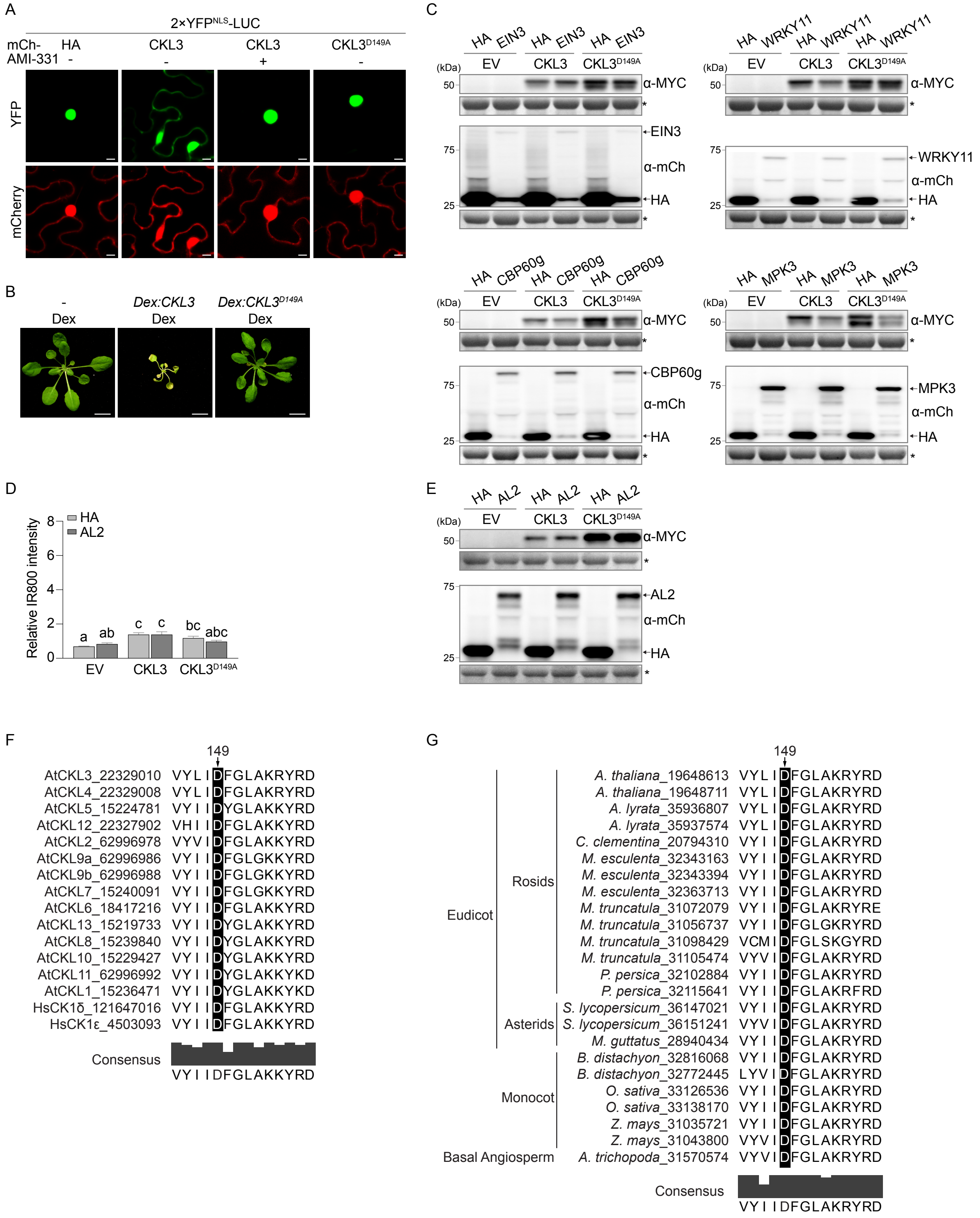
A
2×YFP^NLS-LUC
mCh- HA CKL3 CKL3 CKL3^D149A
AMI-331 - - + -
YFP
mCherry
B
- Dex
Dex:CKL3 Dex
Dex:CKL3^D149A Dex
C
HA EIN3 HA EIN3 HA EIN3
EV CKL3 CKL3^D149A
(kDa)
50
75
25
α-MYC
EIN3
α-mCh
HA
HA WRKY11 HA WRKY11 HA WRKY11
EV CKL3 CKL3^D149A
WRKY11
HA CBP60g HA CBP60g HA CBP60g
CBP60g
HA MPK3 HA MPK3 HA MPK3
MPK3
D
HA
AL2
Relative IR800 intensity
8
6
4
2
0
a ab c c bc abc
EV CKL3 CKL3^D149A
E
HA AL2 HA AL2 HA AL2
EV CKL3 CKL3^D149A
α-MYC
AL2
α-mCh
HA
F
149
AtCKL3_22329010 VYLIDFGLAKRYRD
AtCKL4_22329008 VYLIDFGLAKRYRD
AtCKL5_15224781 VYIIDYGLAKKYRD
AtCKL12_22327902 VHIIDFGLAKKYRD
AtCKL2_62996978 VYVIDFGLAKKYRD
AtCKL9a_62996986 VYIIDFGLGKKYRD
AtCKL9b_62996988 VYIIDFGLGKKYRD
AtCKL7_15240091 VYIIDFGLGKKYRD
AtCKL6_18417216 VYIIDFGLAKKYRD
AtCKL13_15219733 VYIIDYGLAKKYRD
AtCKL8_15239840 VYIIDYGLAKKYRD
AtCKL10_15229427 VYIIDYGLAKKYRD
AtCKL11_62996992 VYIIDYGLAKKYKD
AtCKL1_15236471 VYIIDYGLAKKYKD
HsCK1δ_121647016 VYIIDFGLAKKYRD
HsCK1ε_4503093 VYIIDFGLAKKYRD
Consensus
VYIIDFGLAKKYRD
G
149
Eudicot
Rosids
A. thaliana_19648613 VYLIDFGLAKRYRD
A. thaliana_19648711 VYLIDFGLAKRYRD
A. lyrata_35936807 VYLIDFGLAKRYRD
A. lyrata_35937574 VYLIDFGLAKRYRD
C. clementina_20794310 VYIIDFGLAKRYRD
M. esculenta_32343163 VYIIDFGLAKRYRD
M. esculenta_32343394 VYIIDFGLAKRYRD
M. esculenta_32363713 VYIIDFGLAKRYRD
M. truncatula_31072079 VYIIDFGLAKRYRE
M. truncatula_31056737 VYIIDFGLGKRYRD
M. truncatula_31098429 VCMIDFGLSKGYRD
M. truncatula_31105474 VYVIDFGLAKRYRD
P. persica_32102884 VYIIDFGLAKRYRD
P. persica_32115641 VYIIDFGLAKRFRD
Asterids
S. lycopersicum_36147021 VYIIDFGLAKRYRD
S. lycopersicum_36151241 VYVIDFGLAKRYRD
M. guttatus_28940434 VYIIDFGLAKRYRD
Monocot
B. distachyon_32816068 VYIIDFGLAKRYRD
B. distachyon_32772445 LYVIDFGLAKRYRD
O. sativa_33126536 VYIIDFGLAKRYRD
O. sativa_33138170 VYIIDFGLAKRYRD
Z. mays_31035721 VYIIDFGLAKRYRD
Z. mays_31043800 VYVIDFGLAKRYRD
Basal Angiosperm
A. trichopoda_31570574 VYVIDFGLAKRYRD
Consensus
VYIIDFGLAKRYRD

**Figure S7. $Ca^{2+}$-binding promotes CKL3-Nup58 association for enhanced defense protein nuclear import and ETI execution. Related to Figure 7.**

(A) Cytosolic retention of 2×$YFP^{NLS}$-LUC upon CKL3 and $CKL3^{D149A}$ overexpression. mCherry-tagged HA, CKL3, or $CKL3^{D149A}$ was co-expressed with 2×$YFP^{NLS}$-LUC in *N. benthamiana* leaves for 24 h, followed by pre-treatment with the CKL inhibitor AMI-331 (1 μM) for 12 h prior to confocal imaging. Scale bar = 10 um.

(B) CKL3 overexpression-induced spontaneous cell death in *Arabidopsis*. Ten-day-old transgenic plants carrying *Dex:CKL3* or *Dex:CKL3$^{D149A}$* were treated daily with a 25 μM Dex solution, and images were taken 18 days later. WT plants without the transgene (-) were similarly treated with Dex as a control. Scale bar = 1 cm.

(C) The effect of co-expressing UP-group cargo proteins on CKL3-mediated, $Ca^{2+}$-dependent cell death. mCherry-tagged HA or cargo proteins (EIN3, WRKY11, CPB60g or MPK3) was co-expressed with CKL3-cYFP-MYC (CKL3), $CKL3^{D149A}$-cYFP-MYC ($CKL3^{D149A}$), or empty vector (EV) in *N. benthamiana* leaves. Protein levels were detected at 36 h post-inoculation using Western blotting with an anti-MYC antibody (α-MYC) or an anti-mCherry antibody (α-mCh).

(D and E) The effect of co-expressing AL2 on CKL3-mediated, $Ca^{2+}$-dependent cell death. mCherry-tagged HA or AL2 was co-expressed with CKL3, $CKL3^{D149A}$ or EV in *N. benthamiana* leaves. Cell death-associated autofluorescence was measured using Infrared Fluorescence imaging (IR800) 7 days post-inoculation with relative IR800 intensity presented as mean ± SEM (D). Protein levels were detected at 36 h post-inoculation using Western blotting with an anti-MYC antibody or an anti-mCherry antibody (E). Different lowercase letters indicate significant differences based on a one-way ANOVA, followed by Tukey's multiple comparisons test ($p<0.05$).

(F) Sequence alignment of CKL family members of *Arabidopsis thaliana* (*At*) and CK1 proteins in *Homo sapiens* (*Hs*) at Asp-149. Arrow indicates conservation of Asp-149 in corresponding proteins.

(G) Sequence alignment of CKL3 homologs at Asp-149 within angiosperm plants. Arrow indicates conservation of Asp-149 in corresponding proteins within the species.

*, non-specific bands stained by Ponceau S.

**KEY RESOURCES TABLE**

| REAGENT or RESOURCE | SOURCE | IDENTIFIER |
|---|---|---|
| **Antibodies** | | |
| Mouse monoclonal anti-Myc (HRP conjugate) | Cell Signaling | Cat. #2040S |
| Mouse monoclonal anti-HA (HRP conjugate) | Cell Signaling | Cat. #2999S |
| Mouse monoclonal anti-mCherry | Chromoteck | Cat. #6G6 |
| Goat polyclonal anti-GST | Cytiva | Cat. #27457701 |
| Mouse monoclonal anti-MBP | Invitrogen | REF MA1-10837 |
| Rabbit polyclonal anti-pS149 of Nup58 | This paper | N/A |
| Rabbit monoclonal Phospho-p44/42 MAPK (Erk1/2) (Thr202/Tyr204) (anti-pMPK3/6) | Cell Signaling | Cat. #4370S |
| Rabbit Phospho-(Ser/Thr) Phe antibody (anti-pS/T) | Cell Signaling | Cat. #9631S |
| Goat anti-mouse IgG H&L (HRP) | Abcam | AB97040 |
| Goat Anti-Rabbit IgG (H + L)-HRP Conjugate | BIO-RAD | Cat. #1706515 |
| Donkey anti-Goat IgG (H+L) Secondary Antibody, HRP | Invitrogen | REF PA1-28664 |
| **Chemical, Peptides, and Recombinant Proteins** | | |
| Biotin | Sigma-Aldrich | Cat. #B4501-1g |
| HRP-conjugated Streptavidin | Thermo Scientific | Cat. #N100 |
| Dexamethasone | Sigma-Aldrich | Cat. #D1756-25mg; CAS: 50-02-2 |
| Dexamethasone-Water Soluble (For treatment in protoplast) | Sigma-Aldrich | Cat. #D2915-25mg; CAS: 50-02-2 |
| AMI-331 | Tokyo Chemical Industry | Cat. #A3352 |
| Lanthanum(III) chloride heptahydrate | Sigma-Aldrich | Cat. #262072-100g |
| L-Glutathione Reduced (GSH) | Sigma-Aldrich | Cat. #G4251-10g |
| Maltose | Sigma-Aldrich | Cat. #63418-100g |
| Adenosine 5'-Triphosphate Disodium Salt (ATP) | Sigma-Aldrich | Cat. #A7699-1g; CAS: 34369-07-8 |
| cOmplete Tablets Mini EDTA-free *EASYpack* | Roche | REF 04 693 159 001 |
| PhosSTOP *EASYpack* | Roche | REF 04 906 837 001 |
| Phos-tag ™ Acrylamide | FUJIFILM Wako | Cat. #304-93521 |
| Monolith Protein Labeling Kit RED-NHS 2nd Generation | NanoTemper | SKU: MO-L011 |
| QuantSeq 3' mRNA-Seq V2 Library Prep Kit FWD with UDI (Set A2, UDI12A 0097-0192) | Lexogen | Cat. #194.96 |
| **Bacterial and Fungal Strains** | | |
| *Agrobacterium tumefaciens*, strain GV3101 | N/A | N/A |

| | | |
|---|---|---|
| Escherichia coli NEB 10β | New England Biolabs Inc. | Cat. #C3019H |
| Escherichia coli BL21 | New England Biolabs Inc. | Cat. #C2527H |
| *Pseudomonas syringae* pv *maculicola* (*Psm*) ES4326/AvrRpt2 | Cao et al.[72] | N/A |
| **Experimental Models: Organisms/Strains** | | |
| *Arabidopsis: Dex:AvrRpt2/WT* | McNellis et al.[67] | N/A |
| *Arabidopsis: Dex:AvrRpt2/rps2* | Gu et al.[24] | N/A |
| *Arabidopsis: YFP-YFP-TurboID* | Kim et al.[36] | N/A |
| *Arabidopsis: nup54 nup58* | Ferrandez-Ayela et al.[68] | N/A |
| *Arabidopsis: rps2* | Mindrinos et al.[69] | N/A |
| *Arabidopsis: amiR-CKL3/4.ckl4* | Tan et al.[55] | N/A |
| *Arabidopsis: ckl3-1 (Salk_016571C)* | ABRC | N/A |
| *Arabidopsis: ckl3-2 (Salk_110494C)* | ABRC | N/A |
| *Arabidopsis: TurboID-3×HA-Nup58/Dex:AvrRpt2/WT* | This paper | N/A |
| *Arabidopsis: HA-Nup58/nup54 nup58* | This paper | N/A |
| *Arabidopsis: HA-Nup58*$^{S149A}$*/nup54 nup58* | This paper | N/A |
| *Arabidopsis: Dex:CKL3-mCherry* | This paper | N/A |
| *Arabidopsis: Dex:CKL3*$^{D149A}$*-mCherry* | This paper | N/A |
| *Arabidopsis: Dex:AvrRpm1/WT* | This paper | N/A |
| *Arabidopsis: GCaMP3*$^{NES}$*/Dex:AvrRpt2* | This paper | N/A |
| *Arabidopsis: GCaMP3*$^{NES}$*/Dex:AvrRpm1* | This paper | N/A |
| **Oligonucleotides** | | |
| Primers see Table S6 | | |
| **Recombinant DNA** | | |
| V5-TurboID-NES-pCDNA3 | Addgene | #107169 |
| GCaMP3 | Addgene | #22692 |
| pEG100/TurboID-3×HA-Nup58 | This paper | N/A |
| pEG100/YFP | Gu et al.[24] | N/A |
| pEG100/2×YFP-LUC | This paper | N/A |
| pEG100/2×YFP$^{NLS}$-LUC | This paper | N/A |
| pEG100/ABI5-mCherry | Gu et al.[24] | N/A |
| pEG100/AL2-mCherry | This paper | N/A |
| pEG100/LNK1-mCherry | This paper | N/A |
| pEG100/RPN10-mCherry | This paper | N/A |
| pEG100/CAMBP25-mCherry | This paper | N/A |
| pEG100/WRKY11-mCherry | This paper | N/A |

| | | |
|---|---|---|
| pEG100/EIN3-mCherry | This paper | N/A |
| pEG100/CBP60g-mCherry | This paper | N/A |
| pEG100/SARD1-mCherry | This paper | N/A |
| pEG100/MPK3-mCherry | This paper | N/A |
| pEG100/HA-Nup58 | This paper | N/A |
| pEG100/HA-Nup58$^{S134D}$ | This paper | N/A |
| pEG100/HA-Nup58$^{S146D}$ | This paper | N/A |
| pEG100/HA-Nup58$^{S147D}$ | This paper | N/A |
| pEG100/HA-Nup58$^{S149D}$ | This paper | N/A |
| pEG100/HA-Nup58$^{S150D}$ | This paper | N/A |
| pEG100/HA-Nup58$^{S149A}$ | This paper | N/A |
| pEG100/nYFP-HA-Nup58 | This paper | N/A |
| pEG100/CKL3-cYFP-MYC | This paper | N/A |
| pEG100/CKL3$^{D149A}$-cYFP-MYC | This paper | N/A |
| pEG100/CKL4-cYFP-MYC | This paper | N/A |
| pEG100/CKL3-mCherry | This paper | N/A |
| pEG100/CKL3$^{D149A}$-mCherry | This paper | N/A |
| pEG100/CKL4-mCherry | This paper | N/A |
| pEG100/nLuc | This paper | N/A |
| pEG100/HA-mCherry | Gu et al.[24] | N/A |
| pEG100/HTB1-mCherry | This paper | N/A |
| pEG100/GCaMP3$^{NES}$ | This paper | N/A |
| pBAV154/CFP-RPS2 | This paper | N/A |
| pBAV154/RPS2-CFP | This paper | N/A |
| pBAV154/RPS2-HA | This paper | N/A |
| pBAV154/RPM1-MYC | This paper | N/A |
| pBAV154/NRG1.DV-MYC | This paper | N/A |
| pBAV154/AvrRpm1-HA | This paper | N/A |
| pBAV154/AvrRpm1.GCaMP3$^{NES}$ | This paper | N/A |
| pETL7/GFP | Wang et al.[26] | N/A |
| pETL7/Nup58 | Wang et al.[26] | N/A |
| pETL7/Nup58$^{S149A}$ | This paper | N/A |
| pGEX-4T-1/GST-CKL3 | Tan et al.[55] | N/A |
| pGEX-4T-1/GST-CKL3$^{D149A}$ | This paper | N/A |
| **Software and Algorithms** | | |
| ImageJ | Schindelin et al.[80] | https://fiji.sc |
| Prism 10 | GraphPad Software | https://www.graphpad.com |

| | | |
|---|---|---|
| Snapgene | Snapgene Software | https://www.snapgene.com |
| MaxQuant_2.0.3.1 | Tyanova et al.[81] | https://maxquant.org/maxquant/ |
| Perseus 1.6.15.0 | Tyanova et al.[82] | https://maxquant.org/perseus/ |
| ProteinProspector | N/A | https://prospector.ucsf.edu/prospector/ |
| UCSF Chimera | Pettersen et al.[83] | https://www.cgl.ucsf.edu/echimera/ |
| R | N/A | N/A |
